# Dual Ligand Cooperation at the Plasma Membrane Drives Transport of Engineered Small Extracellular Vesicles Across Brain Endothelial Cells

**DOI:** 10.64898/2026.01.21.700773

**Authors:** Inês Albino, Elena Ambrosetti, Ana Teixeira, Paula Sampaio, Miguel M. Lino, Lino Ferreira

## Abstract

The natural delivery properties of small extracellular vesicles (sEVs) can be harnessed and enhanced through engineering to create a new class of biotherapeutics, particularly for central nervous system (CNS) disorders. While evidence supports the ability of sEVs to cross biological barriers and deliver functional cargo to target cells, a limited understanding of their uptake and transport across the brain hinders their translational potential. In this study, we investigated either native and engineered sEVs, developed by us, using a novel modular engineering platform that employs a dual-targeting strategy to facilitate uptake and transport through human brain endothelial cells (BECs). By utilizing super-resolution microscopy, we provided direct insights into the mechanisms of docking, intracellular sorting, and transport of engineered sEVs. The engineered sEVs formulation demonstrated significantly enhanced uptake, intracellular trafficking across BECs, and the ability to bypass degradative pathways. In vivo, the engineered sEVs exhibited preferential accumulation in the brain choroid plexus, a structure located within the lateral and fourth ventricles, thereby effectively targeting the blood-cerebrospinal fluid (CSF) barrier. These findings highlight the potential of combining advanced targeting strategies with high-resolution imaging to study sEV interactions with the brain biological barriers and develop more effective CNS therapies.

## INTRODUCTION

Small extracellular vesicles (sEVs) are biological nanoparticles of great interest at both fundamental and translational levels[1,2]. They play key roles in cell-cell communication under physiological and pathological conditions and hold potential as vehicles for delivering biomolecules to treat brain diseases[3]. sEVs are recognized for their ability to cross biological barriers at tissue[4–6], cellular[7,8] and intracellular[8,9] levels, facilitating intercellular communication[10,11] and enhancing the therapeutic delivery of biomolecules[5,6,12]. In particular, sEVs derived from mesenchymal stem cells and neural stem cells have demonstrated promising therapeutic effects in various preclinical models of brain diseases, including protective and regenerative outcomes at both local and systemic levels[13–15]. However, their therapeutic application is hindered by extremely low levels of brain accumulation following systemic delivery, typically below 1%[16,17]. This limitation also affects their potential as drug delivery vehicles for transporting exogenous biomolecules into the brain. A major obstacle in improving brain accumulation of sEVs is our limited understanding of how they cross the brain biological barriers including the blood-brain barrier (BBB), the blood-CSF barrier, among others[18]. While a few studies have explored sEV transport across the BBB[19,20] and blood-CSF barrier[11], significant gaps remain[19,20]. For instance, cancer cell-derived sEVs have been shown to interfere with the endocytic pathway in brain endothelial cells (BECs) to avoid degradation and facilitate their transcellular transport[19]. In contrast, little is known about how non-cancer-derived sEVs interact with and cross BECs, or how their transport can be optimized.

To potentiate the accumulation of sEVs in the brain, researchers have engineered the surface of these nanoparticles to improve targeting efficiency[5,21,22] and extend circulation time[23,24]. This has been achieved by modifying the sEVs surface with short peptides, either through post-isolation techniques[22,25,26] or by genetically engineering the sEV-secreting cells[6,21]. Receptor-mediated transcytosis (RMT) has been the preferred pathway for increasing drug delivery across the BBB, utilizing various carriers like antibodies[27], liposomes[28] or sEVs[29]. These carriers exploit the ability of ferrying molecules to bind to receptors and facilitate the transport of drugs into the brain[30]. Studies using single-targeted approaches, where the surface of sEVs is conjugated with a single ligand type to interact with a specific receptor in BECs, have demonstrated improved sEVs retention[25] and accumulation[22] at target tissues or lesion sites compared to non-targeted controls. The efficacy of these strategies has been supported by therapeutic outcomes in preclinical models of stroke, Alzheimer’s and Parkinson’s disease[21,22,25,31]. Furthermore, dual-targeting designs, which involve conjugating the surface of sEVs with two types of ligands, targeting both the RMT receptor and another protein associated with a specific cell type or disease, have been studied to enhance selectivity and brain drug uptake[32,33]. For instance, a recent study explored the use of RGD and Angiopep-2 peptides to increase sEVs accumulation in the ischemic brain[34]. Despite these advances, no study has yet fully investigated the transport mechanisms of modified sEVs or explored the possibility of enhancing their accumulation by combining targeting strategies with transport.

In this study, we investigated the potential to combine sEV targeting strategies with RMT to enhance the crossing of sEVs through BECs. We explored the cooperation between two ligands: hyaluronic acid (HA), polysaccharide that targets the CD44 receptor, recently implicated in endothelial transcytosis[35], and the extensively studied RMT complex, Transferrin-Transferrin receptor (TfR). To increase the spatial proximity between the two ligands, acrylated HA was conjugated to the surface of sEVs, and the remaining vinyl groups were used to attach the RMT ligand via modular “click” chemistry. The human brain endothelial cell line hCMEC/D3 was used in combination with conventional and super-resolution microscopy to provide direct insights into sEVs internalization and transcytosis. Brain accumulation in vivo was evaluated following intravenous administration of native and engineered sEVs in wild-type mice. The biodistribution of sEVs in the brain and main peripheral organs was assessed by ex vivo analysis, and local accumulation was evaluated through immunohistochemical staining. Our results demonstrated that the engineered surface dictates the route of entry and the accumulation of sEVs at the basolateral level for exocytosis, while simultaneously acknowledging the innate ability of native sEVs to escape the degradation pathway. Moreover, our in vivo results indicate that the surface-engineered sEVs accumulated in the brain choroid plexus (blood-cerebrospinal fluid barrier), one of the brain regions with the highest levels of TfR expression[36]. Our findings provide a better understanding of the molecular interactions of engineered sEVs formulations with and through human BECs, bring forth new clues for better targeted designs and spark the discussion about the ability of sEVs to cross brain biological barriers.

## METHODS

### Isolation of plasma from umbilical cord blood

All human umbilical cord blood samples were collected upon signed informed consent, in compliance with Portuguese legislation. The study was approved by the ethical committees of Centro Hospitalar e Universitário de Coimbra (HUC-01-11) and Centro Hospitalar do Baixo Vouga (Ref: 12-02-2018), Portugal. The samples were stored and transported to the laboratory in sterile blood bags with anticoagulant citrate phosphate dextrose adenine solution and processed within 48 h after collection. Briefly, whole blood was diluted 1:1 in EDTA for better separation results, layered on top of Lymphoprep™ (StemCell Technologies SARL, France) and centrifuged at 400 × g for 35 min at room temperature with break off. The upper plasma layer was collected, and the samples were fresh-frozen at −80°C.

### Isolation of plasma-derived sEVs

To isolate plasma-derived sEVs, frozen plasma was allowed to thaw and then centrifuged at 2,000 × g for 20 min at 4°C to pellet dead cells. sEVs were purified by differential ultracentrifugation as described previously[37]. Briefly, samples were ultracentrifuged twice at 10,000 × g for 30 min at 4°C, the pellet was discarded, and the supernatant was submitted to an ultracentrifugation at 100,000 × g for 2 h at 4°C to pellet sEVs. sEVs were then washed with cold PBS and run through commercially available size-exclusion chromatography columns (qEVoriginal 35 nm, IZON Science, New Zealand) in order to separate sEVs from soluble proteins. The sEVs fraction was collected and centrifuged one last time at 100,000 × g for 2 h at 4 °C, resuspended in 150-200 μL of cold sterile PBS and stored at −80°C. All relevant data of our experiment were submitted to the EV-TRACK knowledgebase (EV TRACK ID: EV240161)[38].

### sEVs characterization by NTA

Size distribution and concentration of sEVs was evaluated through NTA using the NanoSight NS300 (Malvern Instruments, U.K.). The system used an O-Ring Top Plate and the sample was injected manually at an approximate flow of 1 mL every 20 s. sEVs were diluted in PBS until a concentration between 15 and 45 particles/frame was reached. For each sample, 5 videos of 30 s were recorded with the camera level set at 15. All the videos were processed with NTA 3.4 analytical software, using a detection threshold between 3 and 5.

### sEVs characterization by electrophoretic light scattering

EV surface charge measurements were performed on a NanoBrook ZetaPALS Potential Analyzer (Brookhaven Instruments Corporation, U.S.A.). Briefly, 8×10^8^ purified sEVs were diluted in 1500 μL of 1 mM KCl solution prepared in biological grade ultrapure water (Fisher Scientific, U.S.A.) filtered twice through a 0.2 μm filter. sEVs were then placed in a disposable polystyrene cuvette and the electrode was immersed within the cuvette. Each sample was measured five times using the Smoluchowski module at room temperature.

### sEVs characterization by total protein quantification

EV protein quantification was performed using the microBCA protein assay kit (Thermo Fisher Scientific, U.S.A.), as per the manufacturer’s instructions. Briefly, bovine serum albumin (BSA) was used to prepare a 9 points standard curve. Then, EV samples were diluted 20 times in 2% (v/v) sodium dodecyl sulphate (SDS) to disrupt the EV membrane. Subsequently, 50 μL of each standard or sample was mixed, in duplicates, with 50 μL of the working reagent provided into a 96-well Corning Costar cell culture plates (CorningInc, U.S.A.). The reaction was incubated for 2 h at 37 °C. Finally, the plates were equilibrated at room temperature for 15 min and the absorbance at 562 nm was read in a microplate reader Synergy™ H1 (Biotek, U.S.A.).

### sEVs characterization: quantification of thiol groups

Quantification of protein thiols was performed using the Thiol fluorescent probe IV (Sigma-Aldrich), as per the manufacturer’s instructions[39]. The fluorescent probe is a coumarin cis-acrylate derivative that is initially non-fluorescent but undergoes a 470-fold fluorescence enhancement upon forming a 1:1 Michael adduct with thiols. It serves as a rapid, highly sensitive, and thiol-specific fluorogenic probe. Briefly, sEVs samples were resuspended in PBS pH 7.4 (1 × 10^10^ particles/mL) and incubated with 10 μM fluorescent probe (1 M stock in DMSO). The presence of DMSO was residual to 0.5% (v/v). The mixture was protected against light and allowed to react for 10 min, and then analyzed with an excitation and emission wavelength of 400 nm and 465 nm using the Synergy H1 microplate reader (Biotek, U.S.A). For each condition, a blank control of the probe in PBS was subtracted to the measured value. The number of thiols was calculated using a standard curve for L-cysteine (Sigma-Aldrich) and normalized to the concentration of sEVs, as estimated by NTA.

### sEVs characterization by Western blot analysis

Western blot analysis for the detection of EV markers and contaminants was performed. Briefly, concentrated EV preparations in PBS (4 μg; approximately 2 to 8 μL) were mixed with 5 × Laemmli buffer (10 μL; 0.35 M Tris-HCl pH 6.8, 10% SDS, 30% glycerol, 0.012% bromophenol blue, 0.6 M DTT) and heated up at 95°C for 5 min. For the analysis of tetraspanins, Laemmli buffer was prepared without reducing agents. The samples were loaded on 12% SDS-PAGE gel (40% bisacrylamide, 1.5 M Tris-HCl pH 8.8, and 10% SDS). The stacking gel was 4% acrylamide containing 40% bisacrylamide, 0.5M Tris-HCl pH 6.8, and 10% SDS. Protein size markers (NZYColour Protein Marker II, NZYTech, Portugal) were loaded without heating. Gel electrophoresis was performed in 1 × Tris/Glycine/SDS buffer prepared from a 10 × concentrated stock (250 mM Tris, 1.92 M glycine, 1% SDS, pH 8.3), initially at 80 V, for 30 min, and increasing to 120 V, for 1 h. Afterwards, PVDF transfer membranes (Amersham Biosciences, Germany) were activated in methanol for 10-15 s and immediately placed in transfer buffer (25 mM Tris, 192 mM glycine, 10% SDS, 20% methanol in water) for 15 min. Then, the gel was stacked on top of the transfer membrane and assembled within a transfer system. The transfer was performed in wet conditions at 100 V for 90 min, on ice. Afterwards, the membrane was removed and blocked in 5% BSA in TBS (2 mM Tris, 15 mM NaCl) with 0.1% Tween 20 (TBS-T) solution for 45 min in a shaker at room temperature. The membranes were then washed with TBS-T and left to incubate overnight at 4°C with the primary antibodies, diluted in 1% BSA/TBS-T: CD63 *1:250* (BD Pharmingen, 556019), ApoA-1 *1:1,000* (Affinity Biosciences, BF0578), GAPDH (G9 clone) *1:250* (Santa Cruz, sc-365062), Calnexin *1:1,000* (Santa Cruz, sc-23954), Alix *1:500* (Santa Cruz, sc-53540), CD9 *1:250* (BD Biosciences 555370), CD81 *1:250* (BD Biosciences, 555675), Hsp70 *1:1,000* (BD Biosciences 554243) and Transferrin *1:1,000* (Santa Cruz, sc-52256). Then, the membranes were washed 3 times with TBS-T and incubated for 1 h at room temperature with polyclonal goat anti-mouse IgG/HRP 1:10,000 (Cell Signaling, 91196) in 1% BSA/TBS-T. The membranes were then washed 3 times and imaged under the VWR® Imager (VWR International, U.S.A.).

### sEVs characterization by Transmission Electron Microscopy (TEM) and immunogold-labelling

TEM analyses of sEVs were performed as previously described[37]. Briefly, sEVs samples were diluted 1:1 in 2% (v/v) paraformaldehyde (PFA) and deposited on Formvar-carbon coated grids (TAAB Technologies, U.S.A.). After washing 4 times with PBS, samples were contrasted with uranyl-acetate 2% for 5 min. sEVs imaging was carried out in a FEI-Tecnai G2 Spirit BioTWIN electron microscope at 100 kV. For immunogold labelling, the grids were floated (sample side facing down) onto 50 μL drops of 50 mM glycine for 3 min to quench free aldehyde groups and then transferred to a drop of blocking buffer (5% BSA/PBS) for 10 min. Then, the grids were incubated in 1% BSA/PBS as a negative control, or anti-Transferrin antibody *1:50* (Santa Cruz Biotechnology, sc-52256) in 1% BSA/PBS for 2 h, followed by six repeated 3 min washes in 0.1% BSA/PBS. The secondary antibody, goat anti-mouse IgC immunogold 15 nm size conjugates *1:200* (TAAB, U.K.) was then incubated for 1 h, followed by additional 2 min washes in PBS, repeated for a total of eight washes. Samples were fixed in glutaraldehyde 1%, for 5 min, and washed in water for 2 min, repeated for a total of eight washes. A final contrasting step was performed using uranyl-oxalate solution pH 7.0 for 5 min, followed by a mixture of 4% uranyl-acetate and 2% methyl-cellulose (1:9) for 10 min on ice. Imaging was obtained using a FEI-Tecnai G2 Spirit BioTWIN electron microscope at 100 kV.

### sEVs membrane labelling

For *in vitro* cellular assays, sEVs were either labelled with the fluorescent dyes Vybrant™ DiO Cell-Labeling Solution (Thermo Fisher Scientific, U.SA.) or PKH67 Green Fluorescent Cell Linker (Sigma-Aldrich, U.S.A.) as per the manufacturer’s instructions. Briefly, the labelling solution (2 µL) was added to concentrated sEVs preparations (100 µL; approximately 1 - 2 × 10^12^ particles/mL) and incubated for 20 min at room temperature. For PKH67 labelling, the EV preparations (2 × 10^12^ particles/mL) were diluted in the kit buffer (diluent C) 1:1, and then PKH67 in diluent C (1:75) was mixed with the diluted sample. Subsequently, samples were incubated for 3 min at room temperature. In both cases, the samples were purified by size-exclusion chromatography, followed by ultracentrifugation at 100,000 × g for 2 h. For stimulated emission depletion (STED) microscopy assays, sEVs were labelled with Atto 647N (Atto 647N DPPE, Atto-Tec GmbH, Germany) suitable for STED imaging. Labelling was achieved by incubating sEVs and Atto 647N (1 mM stock in DMSO) at a molar ratio of 1:20,000 (sEVs:Atto; for 1.3 × 10^11^ particles/mL a final concentration of 13.4 μM Atto 647N was used) in PBS for 2 h in a shaker at room temperature. The presence of DMSO was residual to 1.3% (v/v). Addition of 0.1% BSA/PBS allowed complexation of BSA with unbound label. Samples were purified by ultracentrifugation at 100,000 × g for 2 h. For *in vivo* assays, sEVs were labelled with a Cy7 DPPE conjugate prepared in house through addition of an amine reactive Cyanine7 (Lumiprobe, Germany) and a commercially available DPPE lipid (Avanti Polar Lipids, U.S.A.). Briefly, sEVs were incubated with Cy7 DPPE (0.99 mM stock in DMSO) at a molar ratio of 1:5,000 (this means that for a concentration of sEVs of 1.5 × 10^12^ particles/mL, a final concentration of 12.5 μM Cy7-DPPE was used) in PBS for 60 min in a shaker at room temperature. The presence of DMSO was residual to 2% (v/v). The reaction was stopped by diluting the sample in 0.1% (v/v) BSA in PBS. The samples were purified by ultracentrifugation as described above. As a control for unbound label precipitation, the same labelling procedures were used in the absence of sEVs. In all cases, the sEVs population was characterized before and after labelling, regarding its size distribution profile and concentration.

### Preparation of acrylated HA

High molecular weight HA (50 mg, 180 kDa, Bioiberica, Spain) was dissolved in deionized water (25 mL) to a final concentration of 0.2% (w/v). The temperature was reduced to 0°C. To the above solution one drop of 0.33 M NaOH was added. The pH changed from 5.4 to 9.5, and the solution was stirred at this pH for 30 min. A mixture of 50 molar equivalents of acryloyl chloride (Merck, Germany) was prepared in dichloromethane (25 mL, Fisher Scientific, UK) and added drop by drop over an hour to the HA reaction mixture. During addition of acryloyl chloride dropwise, the pH was maintained between 8-9 by dropwise addition of 1 M NaOH. After complete addition of acryloyl chloride, the solution was allowed to stir for another hour. Throughout the reaction, low temperature (0-5°C) was maintained. Finally, the reaction mixture was filtered, the filtrate was precipitated using a large excess of cold ethanol (500 mL - 1000 mL) and washed twice with ethanol. It was centrifuged, dialyzed (molecular weight cutoff 12 kDa, Sigma-Aldrich) against deionized water for 72 h and lyophilized. The acrylated polymer was stored at 4°C until further use.

### Characterization of acrylated HA by NMR spectroscopy

For ^1^H-NMR spectroscopic analysis of HA final structure, approximately 5 mg of lyophilized sample was dissolved in 600 µL of deuterated solvent (Merck, Germany). The ^1^H-NMR spectra were recorded in D2O using a pulse of 90° and relaxation delay of 4.0 s on a Bruker Avance III™ instrument at 400 MHz. The residual solvent (non-deuterated) was used as the internal reference. All spectra were processed after a manual phase correction and manual baseline correction to obtain a similar baseline for all the peaks. The integral areas of the terminal vinyl group proton peaks (δ5.5 ppm and δ6.5 ppm) were normalized to the integral area of the methyl proton peak (δ2.0 ppm) of HA by integration with MestReNova software.

### Labeling of acrylated HA

HA-acrylate (HA-A, 0.1 g, DS=30%) was dissolved in 0.2 M HEPES buffer pH 8.0 (20 mL) to obtain a final 0.5% (w/v) polymer solution. FAM-Thiol (BioActs, South Korea) was then added to the reaction mixture at a final concentration of 0.24 mM, achieving a labeling efficiency of 2% of the acrylate groups, assuming a reaction efficiency of 100%. The reaction mixture was stirred for 2 h at a room temperature, protected from direct light exposure. The resulting conjugate was purified in a dialysis membrane (molecular weight cutoff 12 kDa) at 4 °C, three times against a pre-cooled 1% (w/v) NaCl (Applichem, Germany) solution and finally against deionized water. After 72 h of dialysis, the FAM-conjugated HA was lyophilized and stored at 4 °C until further use.

### Isolation of low-density lipoprotein (LDL) from human plasma

LDL was isolated from fresh plasma samples obtained from umbilical cord blood. Briefly, plasma was centrifuged at 100,000 × g for 10 min to eliminate chylomicrons by floatation. Then, OptiPrep™ (StemCell Technologies, Canada) was diluted 5 × in the chylomicron-free plasma to a final concentration of 12% (w/v) iodixanol. HEPES saline solution pH 7.4 was added on top to avoid contaminations. Finally, the plasma was ultracentrifuged at 350,000 × g for 3 h at 16°C, using slow acceleration and deceleration. Approximately 14 fractions of 0.5 mL each were collected and subsequently characterized by a SDS-PAGE gel. The fractions enriched in ApoB lipoproteins (>200 kDa) were pooled, concentrated using 3 kDa Amicon filters (3 kDa, PALL, U.S.A.) and used fresh after isolation.

### Apo-Transferrin, LDL and CD98hc labelling

Apo-Transferrin from mouse and human origin (Sigma-Aldrich), LDL and the CD98hc antibody (Invitrogen, MA1-19195) were either labelled with DyLight 550 NHS-Ester or Dylight 650 NHS-Ester (Thermo Fisher Scientific) as per the manufacturer’s instructions. Briefly, the proteins were dissolved in PBS pH 7.4 to a final concentration of 0.5-2.0 mg/mL. A molar-fold excess ratio of 2-7 of reagent was transferred to the reaction tube containing the protein, mixed well and incubated at room temperature for 1 h. Non-reacted reagent was removed from the protein solution by filtration, using Amicon filters (3 kDa) for several washes until the filtrate was clear. The labeled protein was stored in 50% (v/v) glycerol protected from light in single-use volumes at −20°C.

**Pre-loading of apo-Transferrin with Fe (III).** Human apo-Transferrin (Sigma-Aldrich) was loaded with iron using a ferric ammonium citrate solution (Sigma-Aldrich) as the Fe (III) source. Briefly, a solution of ferric ammonium citrate (1 mg/mL) was freshly prepared in 10 mM NaHCO3 and 20 mM HEPES acid, adjusted to pH 7.7. Apo-Transferrin was incubated with the iron solution at 37°C for 10 min to facilitate iron binding. Unbound iron was removed by repeated centrifugation and washing using Amicon filters (3 kDa), until the filtrate was clear. The efficiency of Fe (III) loading was assessed by measuring absorbance at 280 and 465 nm in a microplate reader. Fe (III)-loaded Transferrin was subsequently filter sterilized with a 0.2 μm syringe filter and stored at 4°C until use.

### Apo-Transferrin thiolation

Thiolation of apo-Transferrin was necessary for bioconjugation with sEVs. Thiol-PEG-COOH (6 mg, 1 kDa, Nanocs, U.S.A.) was dissolved in PBS pH 7.4 (30 mL) to obtain a final concentration of 0.2 mM. Afterwards, EDC and NHS (both from Sigma-Aldrich) were added to the reaction at a final concentration of 2 mM and 5 mM, respectively (10:25:1 molar equivalents of EDC:NHS:PEG). The reaction mixture was stirred for 20 min at room temperature. 2-Mercaptoethanol (Sigma-Aldrich) was added at a final concentration of 20 mM to quench the EDC. Then, apo-Transferrin (48 mg, 10:1 PEG:protein) was added and the reaction proceeded for 2 h at room temperature. For the last one hour, rhodamine isothiocyanate (Sigma-Aldrich) was added to the reaction at a final molar ratio of 10:1 (rhodamine isothiocynate:protein). The resulting conjugate was dialyzed (molecular weight cutoff 12 kDa) at 4 °C against PBS pH 7.4. After 48 h of dialysis, the thiol-conjugated protein was lyophilized and stored at 4°C until further use.

### Bioconjugation of sEVs with thiolated ligands

In step 1, FAM-labelled acrylated HA (HA-A-FAM) was dissolved in PBS pH 7.4 to obtain a final 0.35% (w/v) polymer solution. Then, sEVs were added to the polymer mixture at a final concentration of 3 × 10^11^ particles/mL. The reaction proceeded for 2.5 h in a shaker at room temperature. sEVs were purified using ExoQuick ^TM^ precipitation solution (System Biosciences, U.S.A.), as per the manufacturer’s instructions. Briefly, sEVs samples were incubated with ExoQuick reagent diluted 1:5 (v/v) in the sample volume for 30 min on ice, centrifuged for 10 min at 13,000 × g, and the fluorescence in the pellet was measured in a Synergy H1 microplate reader (Biotek, U.S.A) (excitation 480 nm, emission 520 nm). As a control, the same protocol was used in the absence of sEVs, and the fluorescence value of the control sample was subtracted to the measured value. The number of conjugated polymers on the surface of sEVs was determined using a calibration curve to extrapolate the concentration of HA-A-FAM normalized to the concentration of particles, as measured in NTA. In step 2, the remaining acrylate groups on the surface of sEVs-HA-A (approximately 80–85%) were utilized to achieve varying densities of protein immobilization. Apo-Transferrin from mouse and human origins (both from Sigma-Aldrich), LDL, and CD98hc antibody (Invitrogen, USA) with thiol groups were added to the sEVs-HA-A suspension at molar ratios of 1:500, 1:1,000, 1:2,000, and 1:4,000 (sEVs:protein). For a particle concentration of 3 × 10^11^ sEVs/mL, the final protein concentration for the 1:500 ratio was 0.25 μM. The reaction proceeded for 2.5 h while shaking at room temperature. To ensure consistency, the same batch of HA-A was used for all ratios in each protein ligand preparation, minimizing variability associated with HA-A immobilization. For the surface-plasmon resonance (SPR) experiments, sEVs-HA-Tf were incubated for 1 h with a 25 × molar excess of ammonium iron (III) citrate (Sigma-Aldrich) to transform the apo-Transferrin in holo-Transferrin[40]. Afterwards, sEVs-HA-Tf and sEVs-HA-LDL were purified by ultracentrifugation at 100,000 × g for 2 h. sEVs-HA-CD98hc were purified by precipitating the conjugated sEVs with ExoQuick^TM^, as described above, since this procedure did not required sample dilution and was therefore better suited to the lower amounts of antibody available. The fluorescence in the pellet was read at *λ*em = 570 nm (Dylight550) or *λ*em = 670 nm (Dylight650) in a microplate reader. The number of molecules immobilized on the surface of sEVs was calculated from the ratio of protein concentration, extrapolated from a calibration curve, and the concentration of particles measured in NTA. The size and zeta potential of modified sEVs were characterized by electrophoretic light scattering and NTA, respectively.

### SPR experiments

A Biacore T200 and a Biacore X100 (both from Cytiva, U.S.A.) instruments were used to perform SPR experiments. HBS-EP+ (Cytiva) was used as running buffer in the immobilization step, whereas PBS (Thermo Fisher Scientific) was used in the binding analysis. Recombinant Human CD44 (11366-CD) and TfR (2474-TR) proteins, both from R&D Systems, were immobilized via amine coupling reaction on a CM5 gold sensor chip (Cytiva). Briefly, CD44 and TfR proteins were dissolved in PBS (500 μg/mL) and diluted to 50 μg/mL in 10 mM Acetate buffer pH 4.0 before injection for 7 min over a surface preactivated for 7 min with EDC/NHS. The 5-10 min activation period ensures efficient coupling by activating the carboxyl groups on the surface, allowing for optimal protein binding[41]. This resulted in a total of 2500 RU CD44 and 5500 RU TfR immobilized, respectively. Excess reactive groups were deactivated with ethanolamine (1 M at pH 8.5). To prepare the reference channel similar procedure was followed but without injecting the protein. Native sEVs and sEVs-HA-Tf preloaded with Fe (III) were injected over the surface at the concentration of 5.0 × 10^10^ particles/mL in running buffer, using an association phase of 540 s and dissociation phase of 400 s in the case of CD44 surfaces, and an association phase of 600 s and a dissociation phase of 400 s on TfR surfaces, at the flow rate of 10 μL/min. As a control, apo-Transferrin preloaded with Fe (III) and holo- (iron bound) Transferrin (R&D Systems) were injected over the surface at the concentration of 400 nM in running buffer, using an association phase of 300 s and a dissociation phase of 180 s, at the flow rate of 30 μL/min.

### Cell culture

The human brain endothelial cell line hCMEC/D3 was cultured at 37°C in 5% CO2, in EBM-2 medium (Lonza, Switzerland), supplemented with fetal bovine serum (5%, Life technologies, U.S.A.), chemically defined lipid concentrate (1/100, Life technologies), ascorbic acid (5 μg/mL, Sigma-Aldrich), HEPES (2.38 mg/mL, Life Technologies), hydrocortisone (0.51 μg/mL, Sigma-Aldrich) and penicillin-streptomycin (1%, Life technologies). Basic fibroblast growth factor (bFGF, 1 ng/mL, Sigma-Aldrich) was added fresh in the culture medium. Flasks were beforehand coated with collagen I (R&D Systems). Cells between passage 31-35 were seeded at a density of 25,000 cells/cm^2^.

### Cell characterization: endothelial and BBB phenotype

For characterization of junctional proteins, hCMEC/D3 cells were cultured onto type I collagen-coated 96 well-plates (Corning Inc, U.S.A) and ibidi µ-Slide 8 wells (Ibidi, Germany), at a density of 50,000 cells/cm^2^, for 6 days before validation of the model. In both cases, the medium was replaced at day 3. Confluent monolayers of cells were fixed in 4% (v/v) PFA (Alfa Aesar, U.S.A.) for 10 min at room temperature. After permeabilizing the cells with 0.2% (v/v) TritonX-100 (Sigma-Aldrich) for 10 min and blocking for 30 min with 3% (w/v) bovine serum albumin (BSA) solution (Sigma-Aldrich), the cells were incubated with primary monoclonal antibodies solution in 1% (w/v) BSA/PBS for 1 h at room temperature, or overnight at 4°C. Cells were immunostained for the endothelial junctional marker VE-Cadherin *1:100* (Santa-Cruz, sc-9989) and for the junction-associated proteins β-catenin *1:300* (Sigma-Aldrich, 05-665), ZO-1 *1:200* (Invitrogen, 61-7300) and JAM-A *1:100* (R&D Systems, AF1103). After washing, the cells were stained with Alexa-fluor^488^ donkey anti-mouse *1:800* and Alexa-fluor^488^ goat anti-rabbit *1:800* secondary antibodies (both from Invitrogen) in 1% (w/v) BSA solution for 1 h in the dark at room temperature. Cell nuclei were counterstained with 4’,6-diamidino-2-phenylindole (DAPI) (Sigma-Aldrich). Image acquisition was performed in a widefield (INCell Analyzer, GE Healthcare) or confocal fluorescence microscopes (LSM 710, Zeiss) at 40 × magnification, and the images were analysed with ImageJ software (NIH, U.S.A.). To evaluate the expression of receptors and transporters on the cell surface, cells were seeded on type-I collagen coated ibidi µ-Slide 8 wells at a density of 50,000 cells/cm^2^, for 6 days, with the medium replaced on day 3. Subsequently, cells were fixed and immunostained as previously described for the HA native ligand CD44 *1:200* (Invitrogen, 701406), TfR *1:100* (Invitrogen, 13-6800), CD98hc *1:100* (Invitrogen, MA1-19195) and LDLR *1:50* (Santa Cruz, sc-18823), followed by incubation with Alexa-fluor^488^ donkey anti-mouse *1:800* and Alexa-fluor^488^ goat anti-rabbit *1:800* secondary antibodies in 1% (w/v) BSA for 1 h in the dark at room temperature. To evaluate the expression of the t-SNARE protein Syntaxin-4 on the basolateral membrane, cells were seeded on type-I collagen coated ibidi µ-Slide 15 wells 3D, at a density of 50,000 cells/cm^2^, for 2 days, fixed and immunostained for Syntaxin-4 *1:100* (R&D Systems, MAB7894), followed by incubation with Alexa-fluor^488^ donkey anti-mouse *1:800* secondary antibody in 1% (w/v) BSA for 1 h in the dark at room temperature. Cell nuclei were counterstained with DAPI and imaged using the Zeiss LSM 710 confocal microscope with a Plan-Apochromat 40×/1.4 oil immersion objective. Image analysis was performed in ImageJ software.

### sEVs internalization assay

For studying the internalization of sEVs by human BECs, the cells were cultured at a high confluency density of 50,000 cells/cm^2^ on 96 well-plates (Corning Inc) for 6 days, and the medium exchanged at day 3. Before incubation with sEVs, Hoescht 33342 solution (1 μg/mL, ICN Pharmaceuticals, Canada) was added to cell culture medium for 10 min at 37 °C to stain cell nuclei. Afterwards, cells were washed with DPBS (Corning Inc) and incubated for 24 h with DiO-labelled sEVs (1.5×10^9^ particles/mL), native and engineered with the different ligands, in EV-depleted medium. Before sEVs incubation, the fluorescence of the sEVs suspension across experimental conditions was analyzed using a Synergy™ H1 fluorimeter, with excitation and emission wavelengths of 480 nm and 520 nm, respectively. Image acquisition was obtained in a high-content fluorescence microscope (INCell Analyzer, GE Healthcare) using a 20 × objective, with excitation wavelength of 405 nm and 488 nm, at various time-points. After 24 h, cells were washed with DPBS, followed by 5 min incubation with Trypan blue [0.004% (w/v)] to quench the fluorescence of non-internalized sEVs and imaged again. All conditions were performed in quadruplicates. For image analysis, we used the image analysis toolbox of INCell Developer software. Per well, nine images (regions of interest) were used to count the total number of nuclei. An expansion (dilation) of the nuclear mask was applied to define the cytoplasm boundary. The total sEVs fluorescence intensity inside the cytoplasm boundaries normalized to the total number of nuclei (normalized fluorescence), was used as a measure of sEVs internalization per cell. An independent experiment was defined as a separate bioconjugation reaction performed using sEVs obtained from 1-2 isolations derived from pooled plasma of 2-4 individual donors.

### Internalization blocking assay

As before, human BECs were seeded on 96 well-plates at a density of 50,000 cells/cm^2^ and the monolayer was maintained for 6 days. Hoescht 33342 staining solution (1 μg/mL) was added to the cell culture medium for 10 min at 37°C, removed and the cells washed with DPBS. Then, cells were incubated with the blocking antibodies anti-TfR (1 μg/mL, R&D Systems, MAB2474) or anti-CD44 (10 μg/mL, Invitrogen, 701406) for 1 h at 37°C. Subsequently, PHK67-labelled sEVs-HA and sEVs-HA-Tf (1.5×10^9^ particles/mL) were co-incubated with the antibodies and the cells for 24 h at 37°C. Image acquisition was obtained in the INCell Analyzer microscope using a 20× objective, and excitation wavelengths of 405 nm and 488 nm. Image analysis was performed in the INCell Developer toolbox, as previously described. As internalization control, cells were exposed to sEVs in the absence of blocking antibodies. The concentrations of antibody inhibition were based in values previously reported in the literature[42,43]. The toxicity elicited by each antibody to human BECs upon 24 h exposure was evaluated by Hoescht/propidium iodide (PI) staining. Briefly, Hoescht and PI (Invitrogen) were co-incubated with the cells at 1:1,000 and 1:2,000 dilution, respectively, cells were incubated for 15 min at 37°C, and images were acquired in the GE Healthcare INCell 2200 Analyzer imaging system using a 20 × objective, with excitation wavelength of 488 nm and 561 nm. Per well, nine different regions of interest were used to count the total number of nucleated cells and total number of dead cells. This was used as a measure of cell viability, calculated by the formula: (total number of cells - number of dead cells)/total number of cells.

### Spatial organization of CD44 and TfR in BECs after sEVs exposure

Human BECs were cultured at a density of 50,000 cells/cm^2^ on ibidi µ-Slides 15 well 3D glass-bottom (Ibidi), until reaching confluency. In the first set of experiments, the molecular rearrangement of CD44 and TfR in the cell membrane upon sEVs internalization was monitored by Airyscan super-resolution microscopy. Cells were incubated with non-labelled sEVs and sEVs-HA-Tf (1.5×10^10^ particles/mL) for 1 h at 37°C, washed with DPBS and fixed with 4% (v/v) PFA, for 10 min. In a different set of experiments, the interactions of sEVs with CD44 and TfR upon docking was followed. Cells were incubated with PKH67-labelled sEVs and sEVs-HA-Tf (1.5×10^10^ particles/mL) for 1 h on ice, washed and fixed with 4% (v/v) PFA, for 10 min. In both cases, the cell monolayer was incubated with 3% BSA blocking solution for 30 min and stained for CD44 *1:200* (Invitrogen, 701406) and TfR *1:100* (Invitrogen, 13-6800) for 1 h at room temperature without permeabilizing the cell, followed by incubation with Alexa-fluor^555^ goat anti-mouse *1:800* (Invitrogen) and Alexa-fluor^633^ goat anti-rabbit *1:800* (Invitrogen) secondary antibodies. Cell nuclei were counterstained with DAPI and imaged in a Zeiss LSM 980 laser scanning confocal microscope. Images were acquired using a Plan-Apochromat DIC 63x/1.4 objective and the Airyscan 2 detector in Super Resolution mode. Z-stacks of all channels were acquired with a 0.13 µm step size. DAPI, PKH67, Alexa-fluor^555^ and Alexa-fluor^647^ fluorescence was detected using 405, 488, 561, 639 nm for excitation, respectively. The detection windows used for each channel were 422-477 nm, 499-548 nm, 573-627 nm and 659-735 nm, respectively. Images were then processed using Airyscan Joint Deconvolution with 20 iterations. For visual display, Z-sections of all channels were summed and projected in the z-dimension (maximal intensity) and merged using the image analysis software ImageJ. Cropping of images and adjustments of brightness were identical for each labeling and done using ImageJ and Adobe Illustrator (Adobe, U.S.A.). All conditions were performed in triplicates. Image analysis was processed in Imaris 10.1 software (Oxford Instruments, U.K.). An independent experiment is defined as a separate bioconjugation reaction performed using sEVs obtained from 2-3 isolations derived from pooled plasma of 4-6 individual donors.

### Imaging sEVs intracellular trafficking with super resolution and confocal microscopy

Human BECs were cultured at a density of 50,000 cells/cm^2^ on glass-bottom microslides (Ibidi, Germany) until reaching confluency. Then, cells were incubated with Atto 647N-labelled sEVs, sEVs-HA and sEVs-HA-Tf (1.5×10^10^ particles/mL) for 2 h at 37°C. Transferrin-594 conjugate (Invitrogen) was incubated at 25 μg/mL for the last 30 min of the assay. Subsequently, confluent monolayers cells were washed and fixed with 4% (v/v) PFA for 10 min at room temperature. For experiments involving fixed sEVs preparations, saponin (50 μg/mL) was used instead of TritonX-100 to permeabilize the cell membrane without disturbing the surface of sEVs. After blocking for 30 min with 3% (w/v) BSA solution, the cells were stained with anti-Syntaxin-4 *1:100* (R&D Systems, MAB7894) solution in 1% (w/v) BSA for 1 h at room temperature, followed by 1 h incubation with goat anti-mouse STAR Orange *1:100* (Abberior, Germany) at room temperature. Cell nuclei were counterstained with DAPI. STED imaging was performed in a single point scanning confocal Stellaris 8, equipped with a fully motorized inverted Leica DMI8 microscope, a HC PL APO CS2 86x/1.20 STED WHITE water immersion objective equipped with a motorized collar correction (Leica Microsystems), white light laser (WLL), Falcon and STED modalities. The STED images were recorded sequentially by frame scanning unidirectionally at 400 Hz using the galvanometer-based imaging mode, with a line accumulation of 16, and a pixel size of 33 nm for an area size of 33.83 μm × 33.83 μm (1024×1024, zoom factor of 4, pixel dwell time of 0.832 μs) in Leica LasX software (version 4.6.1). DAPI fluorescence was captured with a 405 nm diode laser line with a laser power of 0.85%. Transferrin (Alexa-fluor^594^) was excited with a 590 nm laser line with a laser power of 1.75%, and emission was collected on a Leica HyDX2 (counting mode) detector with a collection window of 600-637 nm. sEVs, labelled with Atto 647N were excited with a 646 nm laser line with a laser power of 6.1%, and emission was collected on a Leica HyDX4 (counting mode) detector with a collection window of 659-755 nm. Syntaxin-4, labeled with Abberior STAR Orange was excited with a 590 nm laser line with a laser power of 1.7 %, and emission was collected on a Leica HyDX2 (counting mode) detector with a collection window of 600-637 nm. The STED depletion was performed with a synchronized pulsed 775 nm (set at 40 %) depletion laser at 30%. For all images the pinhole size was 152.1 μm, calculated at 1 AU for 580 nm emission. The bottom most plane of the glass slide was identified by imaging in confocal reflection mode. Image analysis was performed in Imaris 10.1 (Oxford Instruments, U.K.). In a different experiment, the cells were incubated with PKH67-labelled sEVs, sEVs-HA and sEVs-HA-Tf (1.5×10^10^ particles/mL) for 2 h at 37°C. As a control experiment, Dylight550-labelled apo-Transferrin (0.2 μM) and Cy3-labelled human serum albumin (HSA) (2 μM, Jackson ImmunoResearch, U.S.A.) were incubated with the cells for 2 h, at 37°C. Then, cells were washed, fixed with 4% (v/v) PFA, permeabilized with saponin (50 μg/mL) and immunostained for Rab11 *1:100* (Invitrogen, 71-5300), followed by incubation with Alexa-fluor^633^ goat anti-rabbit *1:800* (Invitrogen). Double and triple immunostainings were visualized in a Zeiss LSM 710 laser scanning confocal microscope. Images were acquired using a Plan-Apochromat 40x/1.4 oil immersion objective and sequential acquisition of separate channels. Image analysis was performed in ImageJ software.

### Image analysis in Imaris

Images were analyzed at the cell’s surface and basolateral level to evaluate sEVs receptor interactions. Z-stack images were three-dimensional (3D)-rendered and analyzed using Imaris software (Bitplane Inc, Switzerland, version 10.1). The size and shape of sEVs and receptors was visualized according to their labellings or immunostainings as a direct map of the intensity distribution of the signal as detected by Imaris. We used “Spots” and “Surfaces” as the creation tools to generate the 3D rendering of complex 3D objects. In addition to that, the smoothing and background subtraction was held constant within a given set of experiments, while the 3D rendering settings for segmentation were adjusted to provide improved 3D rendering upon visual inspection. Any adjustments to the rendered surface shape were made from the edit tool ‘cut surface’ to split surfaces merged during detection, rather than manually shrink or expand the 3D model. For measurements between receptor markers and sEVs, the statistics “Shortest distance to surfaces” was calculated and extracted from Imaris software.

### Animals

C57BL/6J mice were obtained from Charles River, were bred and maintained in the UC-Biotech animal facility, at the University of Coimbra. The animals were housed in groups of 3 to 4 per cage, with food and water provided *ad libitum* and maintained in a 12 h light/dark cycle in temperature and humidity-controlled rooms. All experiments were carried with the approval of the animal ethics committee of the Center for Neuroscience and Cell Biology, University of Coimbra (Ref: ORBEA 262/2020), the approval of the Portuguese DGAV (Ref: 2020-06-29 009414), and in accordance with EU directives regarding animal use in research. No randomization was used to allocate animals to experimental groups.

### *In vivo* experiments

C57BL/6J female mice were subjected to intravenous administrations of DPBS, sEVs or sEVs-HA-Tf. Briefly, the animals were anesthetized with 2.5% isoflurane, and then, in the supine position, were intravenously administered into the lateral tail vein using a 1 mL insulin syringe. The animals were administered with approximately 150 μL of Cy7 DPPE-labelled sEVs and sEVs-HA-Tf suspension (7.5×10^10^ particles/mL). One hour after administration, the mice were anesthetized with isoflurane and were transcardially perfused with 20 mL of ice-cold DPBS to remove residual blood from the brain vasculature. For biodistribution studies, the brain and peripheral organs were carefully extracted and immediately imaged using an IVIS Lumina XR equipment (Perkin Elmer, U.S.A.). The imaging parameters were as follows: λex745 nm, λem 810-875 nm, fluorescence exposure time set to automatic, medium binning, f/stop=2 and a field of view of 10 cm. For fluorescence quantification studies, after perfusion of cold-DPBS, the mice were subsequently perfused with 20 mL of 10% formalin. The brain was carefully extracted and immediately placed in ice-cold 10% formalin for 24 h post-fixation at 4°C. In a different set of experiments, the brain was extracted and immediately embedded in O.C.T. compound (VWR) in dry-ice before being transferred for −80°C for storage.

### Characterization of brain samples by immunofluorescence

The brain was harvested and fixed in 4% (w/v) PFA for 24 h at 4°C, then cryo-protected in 30% (w/v) sucrose in DPBS and allowed to settle for an additional 48 h at 4°C. For Cryostat sectioning brains were snap-frozen before cutting. Next, the brains were embedded in O.C.T. and mounted onto a cryostat (Leica) for sectioning. The brains were serially sectioned with a thickness of 50 μm resulting in five series of sagittal sections covering the entire brain. The sections were kept at −80°C. For the immunofluorescence procedure, sections were treated with blocking solution (5% BSA in DPBS) for 2 h and incubated with the primary antibody (in blocking solution) overnight at 4 °C. Sections were washed with PBS and incubated in the secondary antibody at room temperature for 1 h. Finally, sections were washed three times in PBS, counterstained with DAPI (1 μg/mL) and mounted on glass slides and precision cover glasses #1.5H Thickness (both from Thermo Fisher Scientific) in a fluorescent mounting medium (Dako, Denmark). For non-fixed brains, O.C.T. frozen tissue blocks were transported in dry ice to a cryostat and mounted in cryostat chucks. The brains were serially sectioned and stored at −80°C until the immunohistochemical procedure. Sections were treated with 50 μL of ice-cold 4% (w/v) PFA immediately upon removal from de freezer, fixed for 10 min, and immunostained as described above. Tile-scan pictures of the sections were either obtained in the Cell Observer widefield microscope (Zeiss) or in the SlideScanner widefield microscope (Zeiss) and stitched using Zen black software. Image analysis and quantification were done using ImageJ.

### Statistical analyses

All the results showed in this work are presented as an average of the number of samples for each condition and the standard error of the mean (SEM). Statistical testing was performed using GraphPad Prism®9.0 software. Statistical significance of the results was determined using a parametric unpaired t-test or an equivalent non-parametric Mann-Whitney test, for data with normal and non-normal distribution, respectively. For multiple comparisons between groups, Tukey’s post hoc test was used with one-way ANOVA and Bonferroni’s post hoc test was used with two-way ANOVA. A *P* value < 0.05 was considered statistically significant.

## RESULTS

### Preparation and characterization of sEVs engineered with ligands to enhance BEC targeting and transcytosis

Human plasma sEVs derived from umbilical cord blood, referred to as native sEVs, were chosen for their relatively high yields after isolation and innate immuno-regulatory and regenerative potential[29]. All sEVs were initially isolated using a standard differential ultracentrifugation protocol followed by a size-exclusion chromatography step. sEVs were then characterized based on the recommended Minimal Information for Studies of Extracellular Vesicles (MISEV) characterization guidelines[44]. To further ensure our preparation was enriched in sEVs, western blot analyses were performed to detect common sEV markers and potential contaminants in one batch of pooled donor plasma-derived sEVs (Supplementary Figure S1a). Our results showed that sEVs expressed typical exosomal markers, including CD63, CD9, CD81, Alix, and HSP70. Calnexin, an endoplasmic reticulum marker, and Apo-A, a contaminant found in high-density lipoproteins, were not detected in the sEVs samples.

Similarly, nanoparticle tracking analyses (NTA) revealed that the majority of sEVs had a size distribution within the 100-200 nm range. Electrophoretic light scattering analyses indicated a zeta potential of -20 mV. Regarding purity, sEVs demonstrated an average of 2.9 × 10⁹ ± 2.6 × 108 particles/µg of protein (mean ± SEM across 30 sEVs isolations), which are aligned with previously reported purity ratios achieved using similar protocols with such complex sources[45]. Furthermore, transmission electron microscopy (TEM) analyses showed the presence of an expected cup-shaped structure, with a typical size between 100-200 nm and morphology consistent with individual sEVs (Supplementary Figure S1b).

Next, sEVs were engineered with two ligands to enhance BEC targeting and transcytosis properties. The first ligand was HA, a polysaccharide that was directly immobilized onto the sEVs surface, also serving as an attachment point (i.e., a linker) for the second ligand, one of three RMT protein ligand candidates (see below). To immobilize the HA to the surface of sEVs, we introduced vinyl groups to the polysaccharide (30-40% substitution, meaning 30-40 vinyl groups per 100 disaccharide units of β4-glucuronic acid-β3-N-acetylglucosamine) and then reacted part of these vinyl groups with the protein thiols of sEVs via thiol-ene addition (Figure 1a and Supplementary Figure S2). sEVs-HA conjugates were purified by precipitation with ExoQuickTM. The surface of unreacted sEVs contained approximately 13,692 ± 4,874 free protein thiol groups per vesicle (mean ± SEM, n=4), as determined by a fluorescent probe assay. Following conjugation with HA, this thiol count decreased by approximately 60% (Figure 1b.1). An average of 289 chains of HA were covalently immobilized on the surface of sEVs as extrapolated from the concentration of fluorescently labelled HA in the pellet (Figure 1b.2). Conjugation with HA caused a negligible increase in the mean size (sEVs: 157 nm; sEVs-HA: 162 nm) and a zeta potential of -29 mV and -30 mV, for native and engineered sEVs, respectively (Figure 1b.3 and Figure 1f). In a subsequent step, the remaining acrylate groups in HA were reacted with variable densities of RMT proteins to enhance transcytosis. For this purpose, we have selected protein ligands against two of the most representative RMT targets, Transferrin receptor (TfR)[46] and low-density lipoprotein receptor (LDLR)[47], and the solute carrier CD98 heavy chain (CD98hc)[48]. All ligands carried at least one available thiol group, either naturally (LDL and CD98hc) or through initial thiolation (apo-Transferrin), were fluorescently labeled and finally reacted with sEVs-HA. To control the avidity of the nanoformulation (defined as the cumulative strength of multiple ligand-receptor interactions), and thus the transcytosis transport[49], sEVs-HA were conjugated with low, medium, and high densities of RMT proteins. After purification of the different engineered sEVs, the number of protein ligands at the surface of sEVs was calculated from the ratio of protein concentration, extrapolated from a calibration curve, and the concentration of particles measured in NTA. sEVs-HA-Tf displayed an average of 37, 58 and 147 ligands per vesicle, corresponding to low, medium and high ligand density (Figure 1c.1). This corresponds to a ratio of approximately 0.5 Tf ligands per HA chain at the highest density, confirming that Tf immobilization occurred at less than one protein ligand per polymer backbone. sEVs-HA-Tf showed a similar size profile and mean size to native sEVs, as evaluated by NTA (Figure 1c.2). Furthermore, Transferrin immobilization was confirmed by TEM after labelling sEVs-HA-Tf with Transferrin antibodies conjugated with gold nanoparticles (Figure 1c.3) and western blotting (Supplementary Figure S1c). sEVs-HA-LDL displayed an average of 20 (low), 36 (medium) and 48 (high) ligands per vesicle (Figure 1d.1). No changes were observed in the size profile and mean size of sEVs-HA-LDL as compared to native sEVs (Figure 1d.2). The formulation sEVs-HA-CD98 displayed an average of 177 (low), 328 (medium) and 1007 (high) ligands per vesicle (Figure 1e.1). Conjugation with CD98hc antibody did not change the size profile or mean size of sEVs (Figure 1e.2). Conjugation with HA or the proteins caused no statistically significant changes in the zeta potential of sEVs (Figure 1f.).

**Figure 1.**
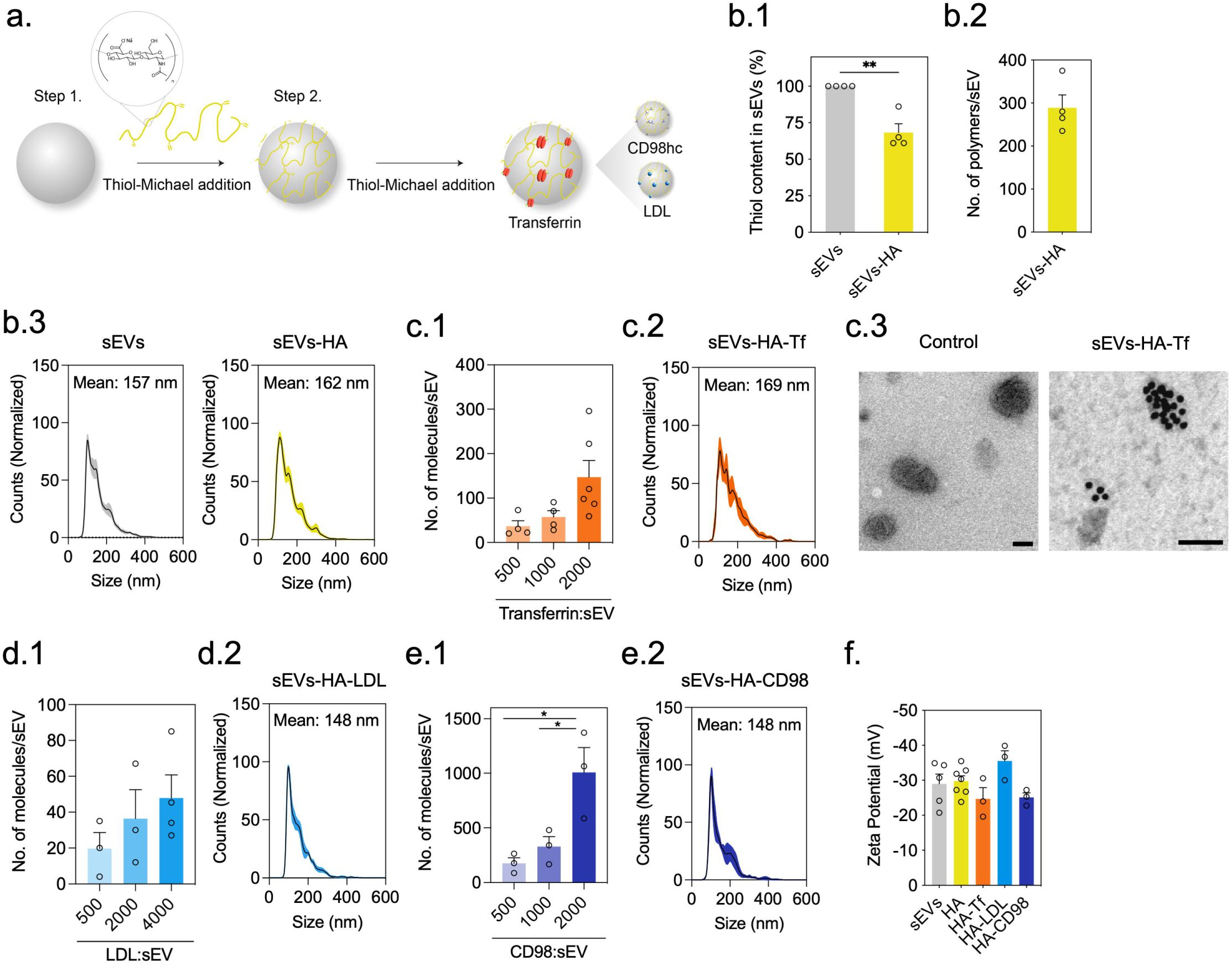
Preparation and characterization of engineered sEVs. (a) Scheme describing the strategy for engineering the surface of sEVs. (b.1) Quantification of available thiol groups on the surface of sEVs before and after conjugation with acrylated HA (HA-A). (b.2) HA-A immobilization on sEVs. The number of immobilized polymers was calculated based on the calibration of the fluorescently labelled polymer, normalized per number of sEVs. In b.1 and b.2, results are expressed as mean ± SEM (n=4 independent sEVs batches and a separate bioconjugation reaction) (b.3) Size distribution profile of sEVs and sEVs-HA evaluated by NTA, expressed in particle counts (percentage) as a function of particle size diameter (nm). Results are mean ± SEM (10 independent experiments). The standard error of the mean is represented in the colour filling. (c.1) Number of Transferrin molecules immobilized on the surface of sEVs at different protein:EV ratios. The number of immobilized proteins was calculated based on the fluorescently labelled protein, as described in the Methods section. The contribution of unspecific binding was evaluated using native sEVs, i.e. before conjugation with HA-A. Results are expressed as mean ± SEM (4-6 independent experiments). (c.2) Size distribution profile of transferrin sEVs conjugates, evaluated by NTA. Results are mean ± SEM (6 independent experiments regarding the highest ligand density tested). (c.3) TEM images of immunogold labelling of Transferrin molecules immobilized on the surface of EVs (15-nm gold particles). Scale bar = 100 nm. (d.1) Number of LDL molecules immobilized on the surface of sEVs at different protein:sEV ratios. Results are expressed as mean ± SEM (3-4 independent experiments) (d.2) Size distribution profile of sEV-HA-LDL, evaluated by NTA. Results are mean ± SEM (6 independent experiments regarding the highest ligand density tested). (e.1) Number of CD98hc antibodies immobilized on the surface of sEVs at different antibody:sEV ratios. Results are expressed as mean±SEM (3 independent experiments) (e.2) Size distribution profile of sEV-HA-CD98. Results are mean±SEM (5 independent experiments regarding the highest ligand density tested). (f) Zeta potential measurements of the sEVs conjugates. Results are mean ± SEM plotted for 3-5 independent experiments. An independent experiment is defined as a separate bioconjugation reaction performed using sEVs obtained from 1-2 isolations derived from pooled plasma of 2-4 individual donors. Statistical analyses were performed by One-way ANOVA followed by Tukey’s multiple comparisons test (p<0.05). In b.1 and e.1 * and ** denote statistical significance (p<0.05 and p<0.01).

Overall, these results indicate that the sEVs surface can be conjugated with controllable amounts of proteins, without significantly changing its surface properties.

### sEVs conjugated with RMT ligands are internalized more efficiently by human BECs than native sEVs

Next, we assessed the level of sEVs internalization by BECs as a surrogate indicator of BEC targeting. The hCMEC/D3 cell line, which has been extensively characterized[50] and widely used for in vitro modelling of BBB processes[51,52], was employed as a model of human BECs. To confirm the BEC phenotype, a confluent monolayer of cells was evaluated for the expression of adherens junctions and tight junctions structural proteins, such as VE-Cadherin, JAM-A, zonula occludens-1 (ZO-1), as well as for β-Catenin, an important Wnt modulator able to induce and stabilize the BBB[53] (Supplementary Figure S3a). Our results showed that the cells expressed most of the key BEC markers. Notably, BECs expressed CD44, the receptor for HA, as well as the TfR, LDLR and the transporter CD98hc, though at variable levels (Supplementary Figure S3b).

The uptake kinetics of sEV formulations conjugated with varying ligand densities by human BEC monolayers were assessed after 6 days in culture. The primary goal was to identify the formulation exhibiting the highest internalization efficiency in BECs. To this end, sEVs were labeled with the fluorescent dyes Vybrant™ DiO or PKH67 and incubated with the cells for up to 24 h (Figure 2a). The intensity of the internalized sEVs signal in the cytoplasm was evaluated at the different timepoints being the fluorescence of non-internalized sEVs quenched with trypan blue. All the surface engineered sEVs formulations showed more BEC internalization than native sEVs. Regarding the surface engineered sEVs formulations, sEVs-HA showed approximately two-times higher internalization by BECs after 18 h incubation when compared to native sEVs (Figure 2b). When sEVs-HA-Tf were used, the formulation conjugated with the highest density of Transferrin significantly outperformed the sEVs-HA formulation at the 12h, 18h and 24h time points (Figure 2c.1 and Figure 2c.2). The formulations conjugated with low and medium densities of Transferrin followed approximately the same internalization kinetics observed for the formulation sEVs-HA. In addition, the sEVs-HA-Tf formulation also exhibited the lowest variability in internalization data among all tested systems. In contrast, no statistically significant differences were observed regarding the internalization of the formulation sEVs-HA-LDL and sEVs-HA-CD98 relative to sEVs-HA, independently of the ligand density (Figure 2d and Figure 2e). However, sEVs with the highest density of ligands (LDL or CD98) showed the highest internalization in BECs, over all the examined densities.

**Figure 2.**
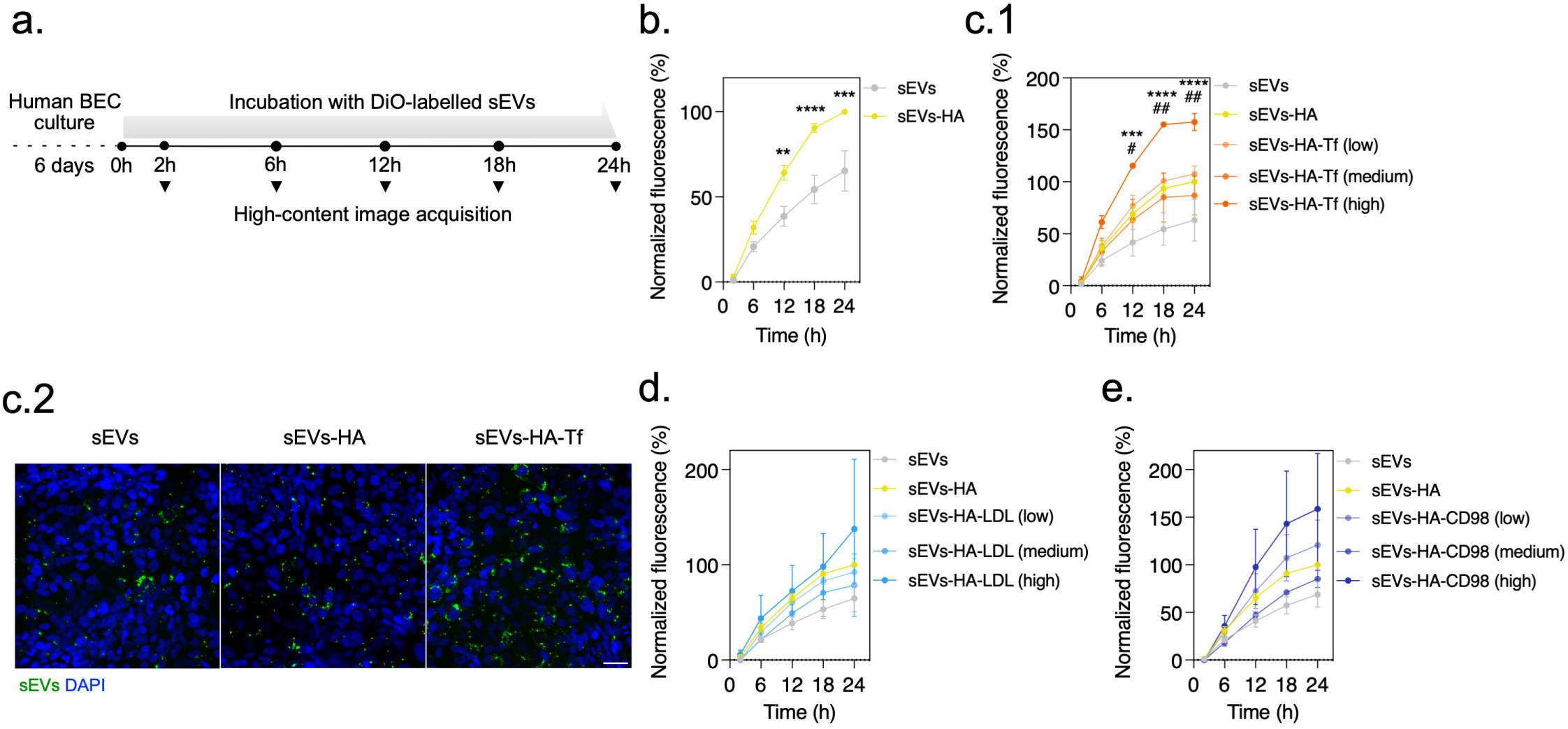
sEVs conjugated with ligands are internalized more efficiently by human BECs than native sEVs. (a) Scheme outlining the workflow of the internalization assay. (b) Internalization of sEVs-HA, following a 24 h kinetics. Fluorescence intensity per cell was normalized to the fluorescence of sEVs-HA at 24 h time-point. Results are expressed as mean ± SEM (n=7 independent experiments; for detailed data analysis refer to the Methods section). An independent experiment is defined as a separate bioconjugation reaction performed using sEVs obtained from 1-2 isolations derived from pooled plasma of 2-4 individual donors. Statistical analysis was performed by Two-way ANOVA followed by Bonferroni’s multiple comparisons test. **, *** and **** denote statistical significance between sEVs and sEVs-HA (p<0.01, p<0.001, p<0.0001). (c.1) Internalization kinetics of sEVs-HA engineered with low, medium and high densities of transferrin. (c.2) Internalization of sEVs-HA-Tf, after 24 h incubation. The observed increase in fluorescence signal corresponded to a higher number of internalized sEV clusters per cell (sEVs: 2, sEVs-HA: 4, sEVs-HA-Tf: 5.4 clusters/cell). Images were acquired in a high content fluorescence microscope. Scale bar: 40 μm. (d-e) Internalization kinetics of sEVs-HA engineered with low, medium and high densities of LDL (d) and CD98 antibody (e). These results were obtained without the need for apo-Transferrin pre-loading, as the concentration of Fe (III) in the culture medium was sufficient to ensure effective saturation of Transferrin with its ligand under the experimental conditions. In c.1, d and e, results are expressed as mean ± SEM (n=3-6 independent experiments). Statistical analysis was performed by Two-way ANOVA followed by Tukey’s multiple comparisons test. # and ## denote statistical significance between sEVs-HA and sEVs-HA-Tf (high) (p<0.05, p<0.01). *** and **** denote statistical significance between sEVs and sEVs-HA-Tf (high) (p<0.001, p<0.0001).

Altogether, our results showed that: (1) sEVs conjugated with ligands were internalized by human BECs more efficiently than native sEVs and (2) sEVs conjugated solely with HA exhibited significant internalization in BECs, which was only surpassed by sEVs-HA-Tf (high). Therefore, for subsequent experiments, sEVs-HA-Tf (high) was selected to study sEVs-BBB interactions.

### sEVs conjugated with HA-Tf interact with CD44 and Tf receptors, and simultaneous inhibition of CD44 and Tf receptors in BECs affects sEVs-HA-Tf cellular internalization

To confirm that the immobilized ligands could interact with the corresponding receptors, we used a surface-plasmon resonance (SPR) assay. In this assay, the sensor chip is coated with the receptor of interest, and the binding capacity of our samples is evaluated by injecting them over the sensor surface (see Methods section). Initial tests were performed with the Transferrin ligand to ensure that these biomolecules could interact with the corresponding receptors, i.e., the TfR. We used both holo-transferrin (iron-saturated) and apo-transferrin (iron-free) pre-loaded with Fe (III) to assess their binding behavior. Since iron binding is known to induce conformational changes in Transferrin that are critical for TfR recognition, the apo-Transferrin was pre-loaded with Fe (III) to restore its receptor-binding capability. The binding response of apo-Transferrin loaded with Fe (III) was similar to that of the holo-Transferrin protein (Supplementary Figure S4), confirming the functional integrity of the ligand-receptor interaction. Then, we evaluated the binding capacity of sEVs and sEVs-HA-Tf to the sensor chip coated with human CD44 or human TfR. As expected, no binding was observed to CD44 or TfR in the case of native sEVs. However, sEVs-HA-Tf pre-loaded with Fe (III) were able to interact with both receptors (Figure 3a.1 and Figure 3a.2).

**Figure 3.**
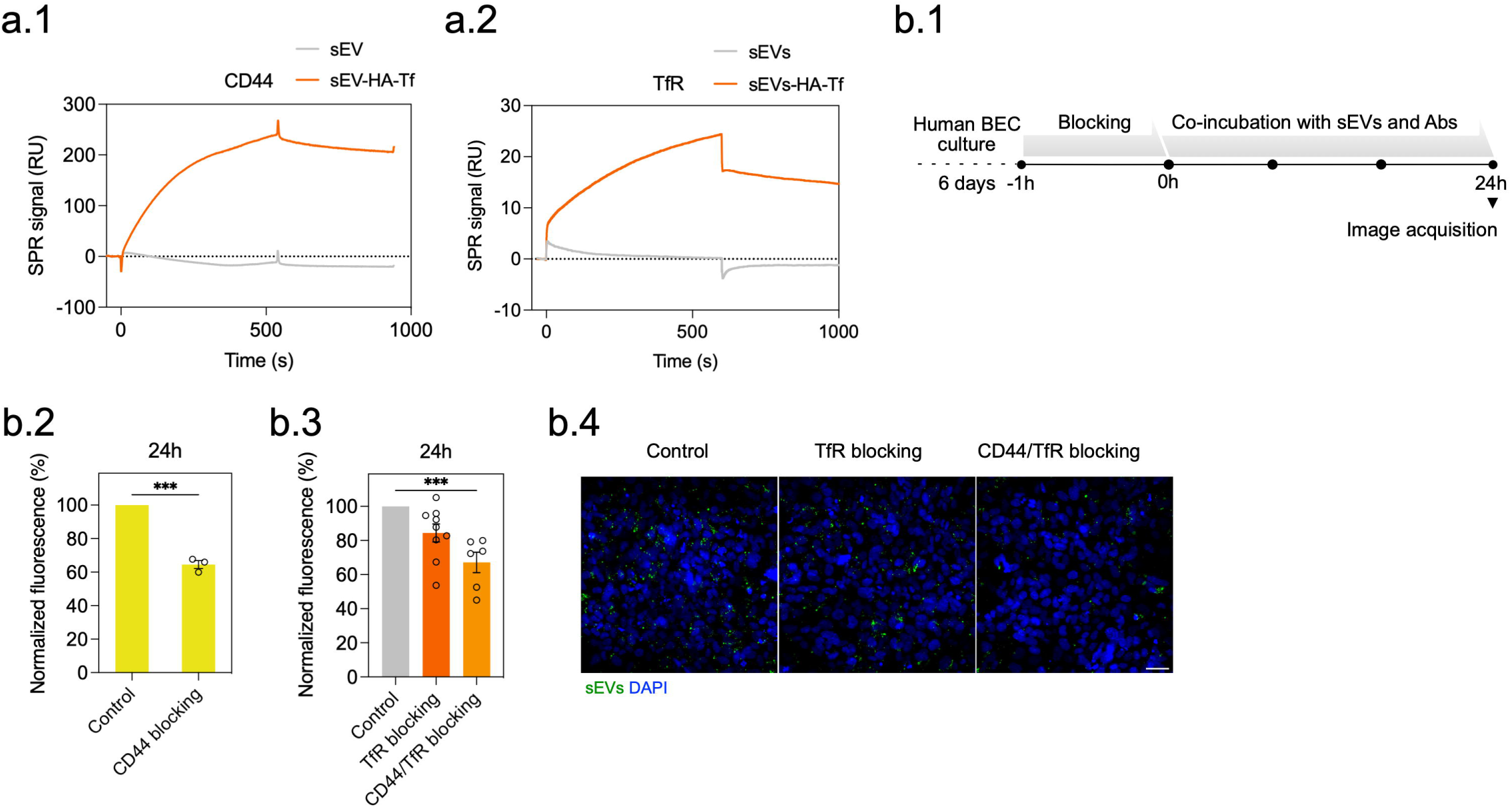
sEVs conjugated with HA-Tf interact with CD44 and Tf receptors. (a.1) Real-time kinetic analysis using SPR for sequential binding of sEVs and sEVs-HA-Tf to the CD44 receptor and (a.2) Transferrin receptor, immobilized on a SPR sensor surface. RU, resonance units. (b.1) Scheme outlining the workflow of the internalization blocking assay. (b.2) Internalization of sEVs-HA, after antibody blocking of the CD44 receptors. Results are expressed as mean±SEM (n=3 independent experiments). (b.3) Internalization of sEVs-HA-Tf, after antibody blocking of the TfR, or CD44 and TfR, simultaneously. Fluorescence intensity per cell was normalized to the fluorescence of the condition sEVs-HA-Tf (control condition, without blocking the receptors). Results are expressed as mean ± SEM (n=6-9 independent experiments). An independent experiment is defined as a separate bioconjugation reaction performed using sEVs obtained from 1-2 isolations derived from pooled plasma of 2-4 individual donors. Statistical analyses in b.2 and b.3 was performed by One-way ANOVA followed by Tukey’s multiple comparisons test. *** denotes statistical significance (p<0.001) between the control condition and the CD44 blocking condition (b.2) or the dual blocking condition (b.3). (b.4) sEVs internalization following 24 h receptor blocking. Images were taken in a high content fluorescence microscope (INCell Analyzer, GE Healthcare). Scale bar: 40 μm.

To confirm that sEVs-HA-Tf internalization was mediated by the interaction with CD44 and/or TfR, we incubated BECs with CD44 and TfR blocking antibodies and evaluated the internalization kinetics of sEVs-HA and sEVs-HA-Tf for 24 h, in the presence of the antibodies (Figure 3b.1). The concentrations of antibody were based in values previously reported in the literature,[42,43] for which no cytotoxic effect in BECs was observed during the period of the assay (Supplementary Figure S5). CD44 receptor blocking caused a 40% decrease in the internalization of sEVs-HA (Fig. 3b.2). In a similar assay, TfR blocking showed to reduce sEVs-HA-Tf internalization in 20% after 24 h. Importantly, this effect was more pronounced (30% decreased internalization) when both CD44 and TfR receptors were simultaneously blocked (Figure 3b.3 and Figure 3b.4).

Overall, our results supported the specificity of the interactions of sEVs conjugated with HA-Tf for CD44 and TfR and suggests a prospective combined role for these two receptors in the targeting and internalization of engineered sEVs.

### CD44 and TfR cooperate in the docking and internalization of sEVs-HA-Tf to the BEC plasma membrane

Spatial reorganization of cell membrane receptors, driven by plasma membrane remodeling, has been shown to play a crucial role in facilitating transcytosis[54,55].

Using Airyscan super-resolution imaging, we evaluated the capacity of sEVs-HA-Tf to modulate the spatial organization of CD44 and TfR upon interaction with BECs. Hence, native sEVs or sEVs-HA-Tf were incubated with a confluent BEC monolayer of cells for 1 h, cells were fixed and the distance between CD44 and its closest TfR neighbor was evaluated. For the sake of the analysis, the distance is represented as a vector, with the initial point set on the external surface of a CD44 receptor and the terminal point set on the closest external surface of the closest TfR. Objects that are entirely separated are given a positive value, but if two objects are intersected then a negative value is given. If two objects touch each other, then the distance is zero (Figure 4a). CD44 rendering was colour-coded according to the distance to the closest TfR. Thus, intersecting surfaces are represented by cool tones and distanced surfaces are represented by warm tones (Figure 4b). As a control, the basal distribution of CD44 and TfR was evaluated in the absence of sEVs. When cells were incubated with native sEVs, CD44 and TfR moved farther apart from each other, as compared with the basal condition. However, in the presence of sEVs-HA-Tf, the average distance between CD44 and TfR in the BEC membrane significantly decreased (approximately 100 nm) (Figure 4b and Figure 4c). These results suggest a cooperation between both receptors for the interaction with sEVs-HA-Tf.

**Figure 4.**
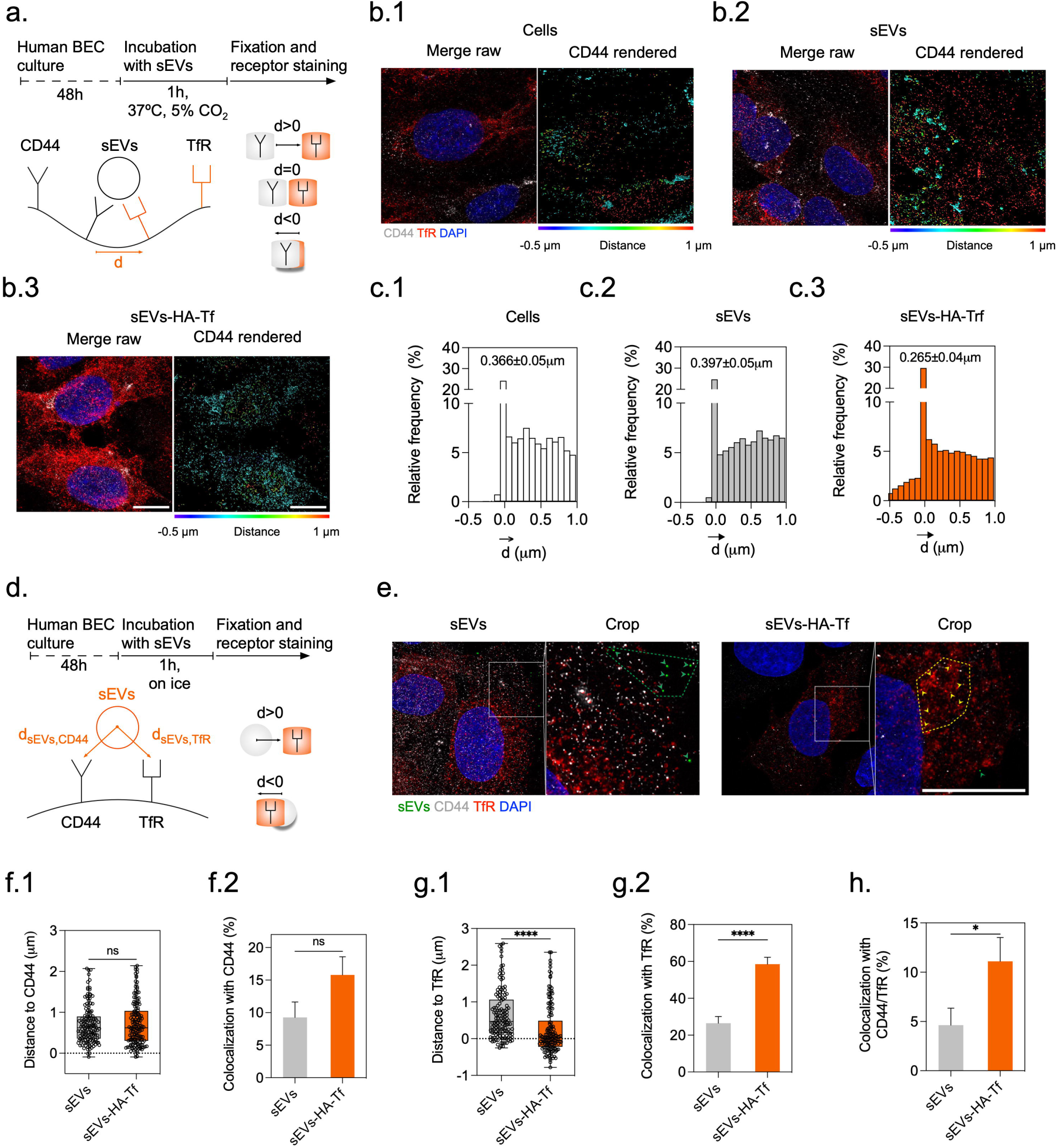
CD44 and TfR cooperate in the docking of sEVs-HA-Tf to the surface of BECs. (a.) Scheme of the experimental setup used to explain the spatial organization of CD44 and TfR at the surface of BECs upon internalization of sEVs. d is expressed as a vector, with initial point set on the external surface of a CD44 receptor and terminal point set on the closest external surface of the closest TfR. (b.) Reconstructed images after deconvolution of CD44 and TfR on the cell surface, upon sEVs exposure, illustrated per condition (b.1) cells alone, (b.2) cells exposed to sEVs and (b.3) cells exposed to sEVs-HA-Tf. CD44 rendered images, illustrated on the right. CD44 rendering is colour-coded according to the distance to the closest TfR. CD44 (gray), TfR (red), DAPI (blue). Scale bar: 10 µm. (c.) Histogram showing the distribution of CD44 with respect to the closest TfR per condition (c.1) cells alone, (c.2) cells exposed to sEVs and (c.3) cells exposed to sEVs-HA-Tf. The values displayed on top of each graph indicate mean±SEM (3643-6173 binned values analyzed per experimental condition across 60-150 cells in 2 independent experiments). (d.) Scheme of the experimental setup used to explain the molecular interactions between sEVs and CD44/TfR receptors upon docking. d is expressed as a vector, with initial point set on the center of the sEVs and terminal point set on the external surface of the closest receptor. (e.) Reconstructed 4-color images after deconvolution and image crops of sEVs docking on CD44 and TfR. Green and yellow arrowheads in the cropped images point to EVs docked on the cell surface. EVs (green), CD44 (gray), TfR (red), DAPI (blue). Scale bar: 10 µm. (f.1) Box plot showing a dot for the distance of each sEVs and sEVs-HA-Tf to the closest CD44 receptor. (f.2) Percentage of co-localization of sEVs and EVs-HA-Tf with CD44. Colocalization is defined as all events below 200 nm. (g.1) Boxplot of the distance of sEVs and EVs-HA-Tf to the closest TfR. Results in f.1 and g.1 are expressed as mean±SEM (136-154 sEVs events analyzed across 40-75 cells in a single experiment). (g.2) Percentage of co-localization of sEVs and sEVs-HA-Tf with TfR. (h.) Percentage of colocalization of sEVs and sEVs-HA-Tf with CD44 and TfR, simultaneously. Results in f.2, g.2 and h. are expressed as mean±SEM (151-171 sEVs events analyzed across 40-75 cells in a single experiment). An independent experiment is defined as a separate bioconjugation reaction performed using sEVs obtained from 2-3 isolation derived from pooled plasma of 4-6 individual donors. Statistical analyses were performed by a Mann–Whitney test (ns: p > 0.05; *: p ≤ 0.05; ****: p ≤ 0.0001). All the data in this panel was acquired in a LSM980 confocal microscope in Airyscan mode and images processed by joint deconvolution.

Then, we analyzed the interactions of native sEVs and sEVs-HA-Tf with the cell surface receptors in conditions where endocytosis was inhibited at 4°C[56]. Native sEVs or sEVs-HA-Tf were incubated with a confluent BEC monolayer of cells for 1 h on ice, fixed and the distance of sEVs towards each receptor separately was evaluated (Figure 4d). Image analyses were performed as above. We classified all events below 200 nm as “colocalization” and the events above 200 nm as “non-colocalization”, which means a surface-surface distance of approximately 100 nm after subtraction of the sEVs radius contribution. We observed that native sEVs predominantly accumulated on a peripheric region, while sEVs-HA-Tf accumulated in close apposition to the nuclei, concomitant with higher TfR expression, as depicted in Figure 4e by the green and yellow arrowheads, respectively. Quantitatively, native and sEVs-HA-Tf showed a similar distribution relatively to CD44, as depicted by the box plot showing a dot for the actual distance of each EV to the closest CD44 (Figure 4f.1). Though, a discreet trend for accumulation of dots at the lower distances was observed in the group sEVs-HA-Tf. This trend was more pronounced in the population under 200 nm, showing higher colocalization of sEVs-HA-Tf with CD44, as compared to native sEVs (Figure 4f.2). Importantly, native and sEVs-HA-Tf showed a statistical different distribution relatively to TfR. Our results showed that sEVs-HA-Tf accumulated within shorter distances to this receptor than native sEVs (Figure 4g.1). The colocalization analysis of sEVs-HA-Tf with TfR was 2.2-fold higher when compared to native sEVs (Figure 4g.2). Finally, we also found 2.4-fold higher colocalization of sEVs-HA-Tf with CD44 and TfR simultaneously, demonstrating preferential binding of the formulation sEVs-HA-Tf to this pair of receptors when compared to native sEVs (Figure 4h).

Taken together, our results suggest cooperation between the two receptors in the docking and internalization of sEVs-HA-Tf at the cell membrane, as indicated by a decreased distance between the molecular entities. Our results further suggest that docking via TfR is more prominent than via CD44.

### sEVs-HA-Tf taken up by BECs accumulate at higher levels in the basolateral membrane than native sEVs

After internalization, most sEVs are moved to early endosomes, where they are sorted into various routes. While sEVs sorted into late endosomes will eventually be transferred to lysosomes for degradation, molecules sorted into recycling endosomes (expressing Rab11[57]) will eventually be transferred both to the apical membrane and to the basolateral membrane for recycling and exocytosis[19,56,58] (Figure 5a). In the basolateral membrane, the SNARE protein Syntaxin-4 regulates the docking and fusion of vesicular compartments with the plasma membrane[59]. To confirm that our cellular model retains some level of apicobasal polarity, we investigated the expression of Syntaxin-4 expression in a confluent monolayer of BECs. Our results indicate that most of the expression of Syntaxin-4 was localized in the basolateral region (between stack 1-3, decreasing until stack 32) of the cell and thus confirming that our model retains a significant degree of apical-basolateral polarity (Figure 5b.1 and Figure 5b.2).

**Figure 5.**
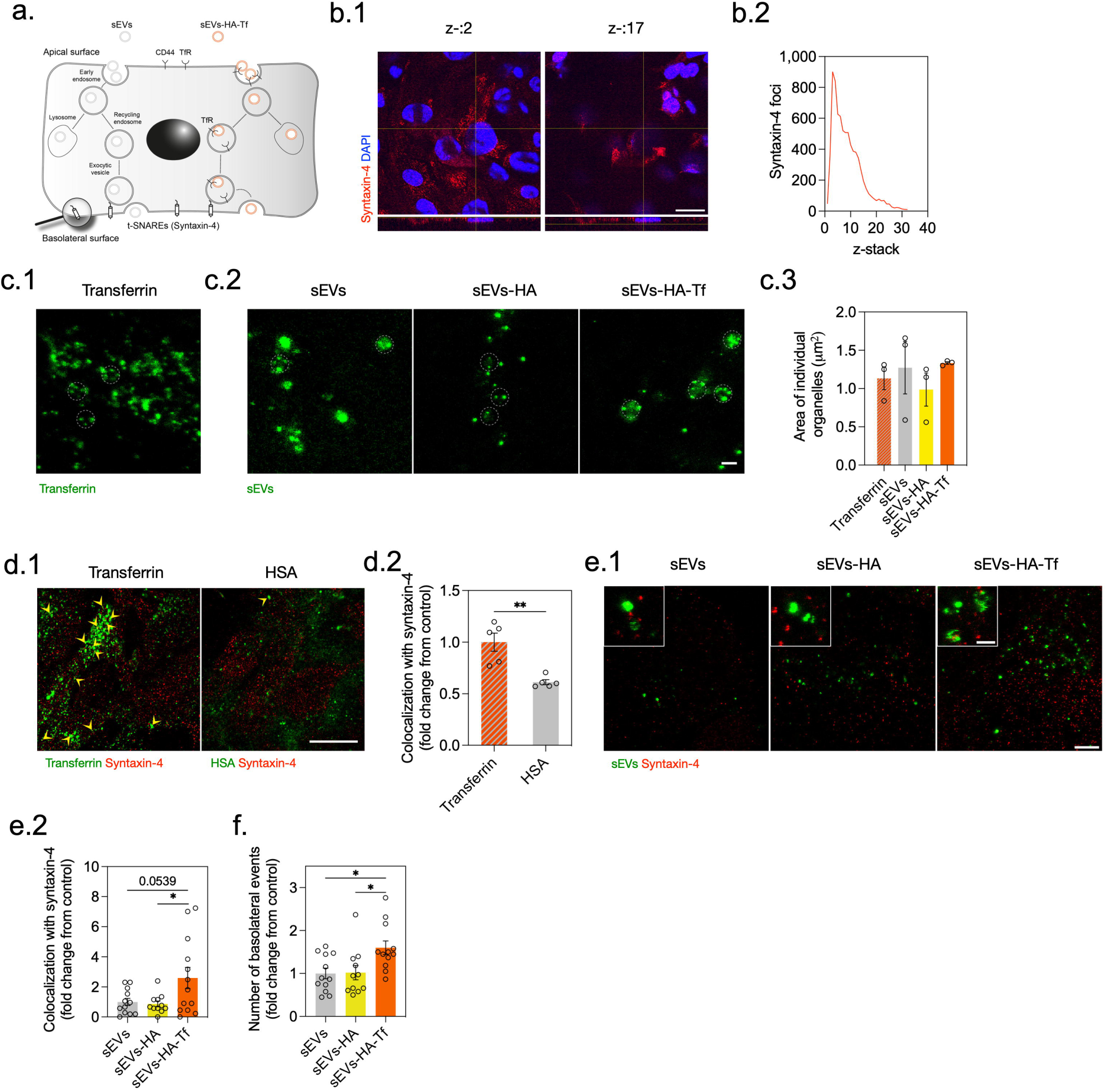
sEVs-HA-Tf accumulate at the basolateral membrane of BECs. (a) Scheme illustrating the trancystosis steps of sEVs or sEVs-HA-Tf in BECs. (b.1) Expression of syntaxin-4 in human BECs along the z-axis. xy-slice at the level of the basolateral (z-stack:2/32) and apical (z-stack:17/32) membranes. Scale bar: 40 µm. (b.2) Number of Syntaxin-4 foci along the z-axis. (c.1) Representative image of Transferrin-594 conjugate, acquired at the bottom most plane of the glass slide, on STED mode using a single point scanning confocal Stellaris 8 (Leica) microscope. (c.2) Representative images of EVs, EVs-HA and EVs-HA-Tf, acquired with similar settings. Scale bar: 1 µm. (c.3) Measured area (μm2) of the encircled organelle structures depicted in c.1 and c.2. Results are expressed as mean±SEM (3 fields analyzed per experimental condition across 105-135 cells in a single experiment). (d.1) Overview of Transferrin and HSA co-localization with syntaxin-4 at the basolateral membrane. Arrowheads point to two-channel overlapping pixels. Scale bar: 20 µm. (d.2) Co-localization of transferrin and HSA with the t-SNARE protein syntaxin-4. Results are expressed as mean±SEM (5 fields analyzed per experimental condition across 175-225 cells in a single experiment). Results are given by the Mander’s co-localization coefficient M1, normalized to the control. Statistical analysis was performed by a Mann–Whitney test (**: p ≤ 0.01). In b.1, b.2, d.1 and d.2, images were acquired in a LSM710 confocal microscope (Zeiss). (e.1) Representative images of sEVs, sEVs-HA, and sEVs-HA-Tf interaction with Syntaxin-4, acquired on STED mode at the plane closest to the glass slide (corresponding to the basolateral side of the cell). Scale bar: 5 µm. Inset: Higher magnification images of EVs in contact with Syntaxin-4. Scale bar: 1 µm. (e.2) Co-localization of sEVs with syntaxin-4, given by the number of sEVs carrying organelles that colocalize with Syntaxin-4. Colocalization is defined as all Syntaxin-4 objects distancing ≤200 nm from sEVs carrying organelles. Results were normalized to the control. (f.) Number of basolateral sEVs carryring organelles, normalized to the control. Results in e.2 and f. are expressed as mean±SEM (11-13 fields analyzed per experimental condition across 25-50 cells in 2 independent experiments). An independent experiment is defined as a separate bioconjugation reaction performed using sEVs obtained from 2-3 isolations derived from pooled plasma of 4-6 individual donors. Statistical analyses were performed by One-way ANOVA, followed by Tukey’s multiple comparisons test (p<0.05).

Next, we attempted to follow the apical-basolateral transport of Transferrin, a well-known example in the transcytosis of iron across BECs[60] and an inspiration for the subsequent sEVs transcytosis study, by a combination of conventional and STED super-resolution microscopy. Human serum albumin (HSA), was used as a negative control, given its comparable size to Transferrin (66 kDa and 76 kDa, respectively), but opposingly lower rate of uptake and transcytosis reported in BECs[61]. Initially, we characterized the levels of colocalization of Transferrin and HSA with the recycling endosome marker Rab11. Transferrin and HSA were incubated with a confluent BEC monolayer for 30 min, on ice, the cells were washed, incubated for 30 min at 37°C, fixed and immunostained for Rab11 (Supplementary Figure 6a). Colocalization with Rab11 was given by the Mander’s colocalization coefficient M1, which measures the fraction of intensity in the protein channel that overlaps with the Rab11 channel, at the pixel level, reflecting spatial correlation within the resolution limits of the imaging system. We observed a higher percentage of internalized Transferrin colocalized with Rab11 compartments (52.8% ± 9.4%), as compared with HSA (34.8% ± 5.4%), despite the percentage of colocalization was not statistically significant (Supplementary Figure S6b.1 and Supplementary Figure S6b.2). Then, we investigated whether native sEVs or sEVs-HA-Tf can proceed via the recycling pathway to transcytosis. We monitored sEVs colocalization with Rab11. sEVs were incubated with a confluent BEC monolayer for 2 h, cells were fixed and immunostained for Rab11 (Figure 2.12a.). Colocalization with Rab11 is given by the Mander’s colocalization coefficient M1, expressing the fraction of intensity in the sEVs channel that overlaps with the Rab11 channel. We observed that all the three groups colocalized with Rab11 in the perinuclear region, but sEVs-HA-Tf (28.8% ± 2.3%) showed higher colocalization than native sEVs (23.1% ± 1.0%) and sEVs-HA (18.9% ± 1.4%) (Supplementary Figure S7).

Next, we focused on exocytosis of native and sEVs-HA-Tf, which occurs when intracellular vesicles fuse with the basolateral membrane of BECs. For this, we used stimulated emission depletion (STED) microscopy to visualize the interaction of sEVs with the basolateral membrane. In our analysis, we defined transporting organelles as large vesicular structures, approximately 1 μm in diameter, that carry sEVs or Transferrin and are involved in the final stages of transcellular transport. We assessed the following parameters: (i) the size of transporting organelles, (ii) the number of exocytotic events at the basolateral membrane, and (iii) their colocalization with the t-SNARE protein Syntaxin-4. Colocalization was determined by the proximity of a transporting organelle to the nearest Syntaxin-4 signal, defined as within 200 nm. A fluorescent Transferrin conjugate was used to study vesicle fusion with the basolateral membrane. For this purpose, Transferrin was incubated with the cells for 30 min, cells were fixed and imaged in the basolateral region (first image of a z-stack pile acquired at the bottom most plane of the glass slide, on reflection mode). We consistently observed fluorescent Transferrin surrounding vesicle-like structures, which seem to be responsible for carrying Tf to the basolateral region of the cell (Figure 5c.1). Interestingly, organelles carrying sEVs or Tf exhibited similar size and fluorescence profiles (Figure 5c.1, Figure 5c.2 and Figure 5c.3). These structures were encircled by fluorescence puncta of approximately 100 nm size, likely representing single or clustered sEVs particles, lining the membrane of the larger 1 µm vesicles. These findings suggest that both Transferrin and sEVs are transported in similar vesicular carriers that ultimately fuse with the basolateral membrane to release their contents.

Next, we quantitatively assessed the colocalization of Transferrin and HSA at the basolateral level with Syntaxin-4 (Figure 5d.1 and Figure 5d.2). We observed a 1.6-fold higher colocalization of Transferrin with Syntaxin-4 compared to HSA, which seems to follow a greater number of Transferrin molecules reaching the basolateral membrane. Together with previous results showing colocalization levels of Transferrin and HSA with the recycling endosome marker Rab11, these results clearly indicate that the Transferrin has a greater chance to be routed to the basolateral membrane for exocytosis, while HSA will be greatly routed to the apical membrane for recycling. We then evaluated the colocalization of native and engineered sEVs with Syntaxin-4. Colocalization is given by the number of sEVs carrying organelles that colocalize with Syntaxin-4. We observed a higher extent of colocalization of sEVs-HA-Tf with Syntaxin-4, that was 2.6-fold and 3.4-fold higher than the colocalization of sEVs and sEVs-HA with Syntaxin-4, respectively (Figure 5e.2). These interactions are depicted in Figure 5e.1 obtained with STED microcopy and exclude the interactions with other proteins besides Syntaxin-4 that are involved in the SNARE complex formation at the basolateral membrane[62]. Finally, adding to previous observations supporting higher colocalization with Syntaxin-4, we have observed a higher total number of sEVs-HA-Tf carrying organelles reaching the basolateral membrane (Figure 5f). These results confirm the hypothesis of facilitated transcytosis of the engineered formulation.

### The intravenous administration of sEVs-HA-Tf led to accumulation in the brain choroid plexuses

Although several studies have shown that the conjugation of nanoparticles with peptides or antibodies against Transferrin improve accumulation in the brain, the levels remain relatively low (typically below 0.1%), due to the highly selective nature of RMT and the anatomical challenges posed by the BBB[63,64]. To study sEVs accumulation and the sites of association to the brain vasculature, we administered a single-dosage of 4×10^11^ particles of Cy7-labelled sEVs and sEVs-HA-Tf in the tail vein of wild-type mice (Figure 6a). Both conditions were derived from the same labeling batch of sEVs, with their concentration and size evaluated prior to administration, confirming no aggregation (Supplementary Figure S8). After 1 h, we immediately imaged the brains ex vivo, or prepared them for later sectioning and immunostaining (Figure 6a). For the latter subset of animals, we either fixed them immediately, after perfusion with a saline solution, or we snap-frozen the brain tissue after perfusion without fixation. We selected a 1 h time point to characterize sEV accumulation in the BBB based on two main reasons: (i) previous studies have shown that approximately 98% of the injected dose is cleared from the bloodstream within 1 h [65,66], and (ii) our own data indicate that brain accumulation does not significantly change beyond the first hour following intravenous administration[67]. IVIS imaging showed that approximately 0.32% of the total signal (sum of radiance normalized per organ weight) of sEVs-HA-Tf accumulated in the brain, compared to 0.21% for native sEVs (Figure 6b.1). The fluorescent signal accumulated in the two conditions is given in a pseudo-color radiance scale in the range of 3.3×107 and 1.2×108 W.cm-2.sr-1 (Figure 6b.2). This represents a 1.5-fold enhancement in brain delivery with sEVs-HA-Tf, in alignment with other systems where Transferrin or Transferrin receptor-targeting ligands are conjugated to nanoparticles to enhance their transport across the BBB, achieving 1.2- to 2.4-fold increases in brain targeting over bare control conditions[68–70]. sEVs distribution in the other main organs was similar in both conditions and comparable to sEVs distributions previously described in literature[16] (Supplementary Figure S9). In parallel, fixed brains were used for staining of the vascular endothelium and imaging of the distribution of the fluorescently labelled sEVs. Interestingly, we observed an accumulation of the sEVs-HA-Tf formulation inside the choroid plexuses, specifically in the lateral and fourth ventricles, when compared to native sEVs (Figure 6c.1 and Figure 6c.2). These observations were supported by higher Transferrin receptor expression along the vasculature of the choroid plexus[71] (Supplementary Figure S10). To avoid artifacts associated with perfusion-fixation processing, we ran a second animal experiment in which the brain of the animals was snap-frozen immediately after perfusion. We observed signal aligned over vessels lining the lateral ventricle in sEVs-HA-Tf treated brains (Figure 6d.1). To quantitatively address these observations, we measured the mean grey value intensity delimited inside the choroid plexus (dashed line) and in its periphery (a ROI defined outside this area), for both native sEVs and sEVs-HA-Tf treated brains (Figure 6d.2). We found increased fluorescence in the choroid plexus in brains treated with the sEVs-HA-Tf compared to the neighboring region (distanced away approximately 500 μm). Instead, brains treated with native sEVs, showed similar grey values both in the choroid plexus and its periphery. Altogether, the sEVs-HA-Tf formulation demonstrated enhanced accumulation in the brain compared to native sEVs. This increase may be attributed to the higher accumulation of sEVs-HA-Tf in the choroid plexus.

**Figure 6.**
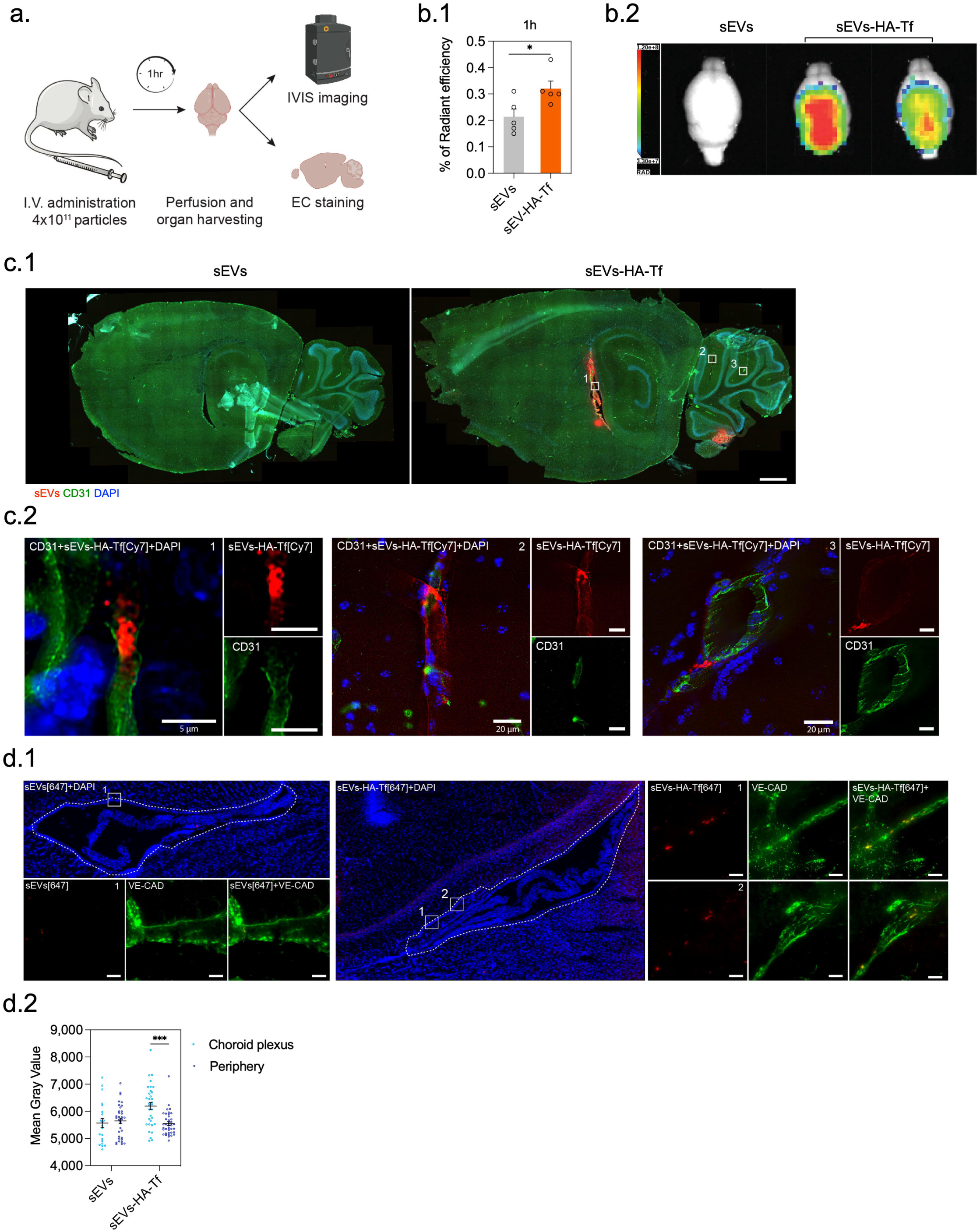
sEVs-HA-Tf accumulate in the brain choroid plexuses. (a.) Schematic drawing of the route and number of administered particles for IVIS imaging and immunostaining. Image icons created with BioRender. (b.1) Quantification of sEVs signal accumulation in the brain after 1 h. Imaging units are defined as radiant efficiency normalized by organ weight. Results are expressed as mean ± SEM (n=5 animals per experimental condition). Statistical analysis was performed by an unpaired t-test. * denotes statistical significance between sEVs and sEVs-HA-Trf (p<0.05). (b.2) EVs signal accumulation in the brain, 1h after administration. Images are presented with a relative photon scale automatically adapted to use the full dynamic range of the pseudocolor scale (3.3×107-1.2×108 W.cm-2.sr-1). (c.1) Representative maximum intensity projections of a large field of view showing brain sagittal sections of sEVs and sEVs-HA-Tf treated brains, acquired in a Zeiss Axio Observer 7 widefield microscope. Images were collected from formalin-fixed brains 1 h after injection of sEVs. Scale bar: 800 µm. (c.2) High-resolution images of the insets showed in c.1. Brain microvasculature immunostained with CD31, acquired with a Plan-Apochromat 40x/air objective, in a Zeiss Axio Observer 7, after deconvolution. CD31 (green), sEVs (red) and DAPI (blue). (d.1) Z-stack projections of a large field of view around the lateral ventricle immunostained with VE-Cadherin and imaged in a slide scanner (AxioScan7) with a 20x objective. Images were collected from non-fixed brains 1 h after injection of EVs. The appended boxes correspond to image crops acquired in the numbered regions. (d.2) Quantification of fluorescence gray values inside the choroid plexus (dashed line) and inside a ROI defined outside this area. Results are expressed as mean±SEM (21-38 foci analyzed per experimental condition across 3-4 sagittal sections). Statistical analysis was performed by Two-way ANOVA followed by Bonferroni’s multiple comparisons test. **** denote statistical significance between the lateral ventricle and the periphery (p<0.0001).

## DISCUSSION

There is a growing consensus that sEVs can cross biological barriers, including tissue, cellular, and intra-cellular barriers, as evidenced by sEVs functional cargo transfer to recipient cells[5,6,12]. However, direct and conclusive evidence of sEVs transport across the brain endothelial barrier remains elusive, largely due to methodological challenges in tracing sEVs and their interactions with subcellular compartments[3,72]. Understanding the mechanisms underlying transcytosis-mediated transport is crucial for enhancing the accumulation of sEVs in the brain, a key objective for therapeutic applications. To address this challenge, we have used super-resolution microscopy to resolve sEVs interactions with both the apical and basolateral membranes of BECs. In addition, we have investigated the transport of sEVs coated with a novel dual-ligand strategy through the brain endothelial barrier and compared it with native sEVs. Our findings provide the first direct demonstration of sEV-mediated simultaneous recruitment of two receptors, CD44 and TfR, revealing a previously unrecognized synergy that facilitates targeted transport across the brain endothelial barrier.

To enhance the performance of sEVs in crossing the BBB, we employed a dual-ligand surface coating leveraging the cooperative interaction between HA and Tf. This strategy not only improved sEV targeting and internalization but also played a critical role in facilitating sEV transcytosis across BECs. This dual-ligand approach builds on earlier strategies that combined engineered neuropeptides and integrin-binding peptides for BBB targeting[21,22].

We first screened various receptors and transporters to identify ligands that would enhance BEC accumulation, using internalization efficiency as a key criterion. Among the tested formulations, sEVs-HA-Tf achieved the highest internalization. By employing endogenous Transferrin, we ensured that the ligand occupies the physiological TfR binding site, initiating a normophysiological endocytosis pathway into BECs. The study did not include a single-ligand sEVs-Tf control because the conjugation chemistry used in our modular design required the initial anchoring of HA to provide reactive acrylate groups for subsequent ligand coupling. Thus, direct Tf attachment was not feasible within the same framework. More importantly, this work is centered on a stepwise, modular design where HA serves both as a functional backbone and as a targeting moiety, onto which an additional RMT ligand such as Tf can be introduced. In this context, our comparisons focused on sEVs, sEVs-HA and the dual-ligand sEVs-HA-Tf to evaluate the incremental effect of combining HA with a RMT ligand. Nevertheless, potential competition with endogenous Transferrin suggests that future efforts should explore alternative ligands or modifications. Furthermore, ligand density is as a critical parameter influencing not only internalization but also exocytosis[73]. While our study focused on internalization, higher ligand densities, although beneficial for uptake, do not necessarily enhance exocytosis. Future studies should further investigate the effect of ligand density in sEV transcytosis.

Our results indicate that the cooperative interaction of CD44 and TfR induced by sEVs-HA-Tf significantly enhanced sEVs docking and internalization in BECs. Super-resolution microscopy revealed that the engineered sEV formulation altered receptor organization, bringing CD44 and TfR into closer proximity at the cell surface. In contrast, native sEVs induced these receptors to remain more spatially separated. Additionally, our engineered formulation docked preferentially to TfR, likely due to a higher number of accessible binding sites and greater spatial availability of Transferrin compared to HA. sEVs-HA-Tf exhibited substantially higher simultaneous colocalization with both CD44 and TfR than native sEVs, underscoring the complementary roles of these receptors. Further supporting this synergy, simultaneously blocking CD44 and TfR significantly reduced the internalization of the engineered formulation, more so than blocking either receptor alone. The partial reduction in uptake observed upon receptor-blocking is in line with previous reports, which have been attributed to incomplete receptor coverage, antibody binding to non-functional epitopes, receptor recycling, or the presence of alternative uptake pathways[42,74]. While the CD44/TfR pair has been briefly explored as a dual-targeting system in imaging probes and drug-conjugated aptamers for tumor or cancer stem cell applications[75,76], it has not been investigated in the context of brain targeting until now. These findings reveal a previously unrecognized synergy between CD44 and TfR and their potential as dual-targeting receptors for sEV-mediated delivery.

As far as we are aware, this is the first study to use super-resolution microscopy to resolve the molecular interactions of engineered sEVs at the plasma membrane and trace their intracellular trafficking. By surpassing the conventional 200 nm diffraction limit and achieving lateral resolutions of 30-100 nm, we gained unprecedented insights into sEV behavior at both the apical and basolateral membranes. Alternative proximity-based methods such as FRET and PLA can detect ligand-receptor proximity (<40 nm) but do not capture the spatial organization or ultrastructural context of these complexes, which super-resolution microscopy provides[77,78]. In the final stages of transcytosis, we observed similar organizational patterns at the basolateral membrane for Transferrin-conjugates and sEVs, suggesting the involvement of analogous transport ultrastructures. While a general mechanism for RMT of TfR-based antibody constructs where transcytosis is driven by sorting along intracellular tubules has been proposed[79], comparable data for sEV transcytosis remain limited, especially for therapeutic formulations.

Existing evidence from metastatic breast cancer cell–derived sEVs in a zebrafish BBB model shows that endocytic vesicles containing sEVs can fuse with t-SNARE proteins such as SNAP23 and Syntaxin-4, lending strong support to the concept of sEV transcytosis[19]. Consistent with this, we observed both native and engineered sEVs (sEVs-HA and sEVs-HA-Tf) localizing at the basolateral membrane and colocalizing with Syntaxin-4. Notably, sEVs-HA-Tf reached the basolateral surface in higher numbers than either native sEVs or sEVs-HA. These observations suggest that TfR interactions promote co-transport while helping bypass degradative pathways. Although other proteins likely contribute to the t-SNARE complex, Syntaxin-4 colocalization provided an adequate readout, enabling us to resolve the final stages of vesicle trafficking with super-resolution microscopy and to gain mechanistic insight into the molecular steps of sEV transport across BECs. In line with our findings, previous studies using in vitro BBB transwell models have shown that only a very small fraction of sEVs (∼0.16%) can be recovered beyond the endothelial barrier, emphasizing that transcytosis is a rare event[19]. Future work may benefit from employing more sensitive quantification methods, such as radiolabeling combined with PET imaging to better capture and quantify these low-frequency transport events.

In vivo, the sEVs-HA-Tf formulation demonstrated significantly higher accumulation in the brains of wild-type mice than native sEVs. Notably, the engineered sEVs, but not native sEVs, preferentially accumulated in the choroid plexus of the lateral and fourth ventricles. Unlike BECs, the endothelial cells of the choroid plexus are fenestrated and lack tight junctions, resulting in a leaky endothelium surrounded by a TfR-rich epithelial layer[36]. These anatomical features likely facilitated the pronounced enrichment we observed at this interface and may contribute to the ability of TfR-targeted nanoformulations, including our engineered sEVs, to access the brain via the blood-CSF barrier. Our engineered sEVs may thus represent a promising therapeutic strategy to address a range of neurological conditions, including neuroinflammatory and infectious diseases, cerebral amyloid angiopathy, brain tumors, and neurohumoral dysregulation[80].

Although CSF-borne substances can reach deep brain structures, the mechanisms for sustained CSF delivery remain poorly understood. Current approaches, such as folate transport[10], sEV release[11], plasma protein transport[81], and receptor-mediated endocytosis[82] offer promising avenues for CNS targeting but require further optimization. By demonstrating the choroid plexus as a viable entry point for sEVs, our study provides a foundation for developing more effective CNS-targeted sEV formulations and expands the potential application of sEV-based therapies to conditions affecting the blood-CSF barrier.

## CONCLUSIONS

The development of new CNS therapeutics is closely tied with the simultaneous development of tailored brain delivery technologies. Identifying the mechanisms by which they operate is needed to understand why many current strategies have failed so far. Here, we demonstrate that engineered sEVs can outperform native sEVs in BEC transcytosis. Importantly, we also demonstrate that native plasma-derived sEVs have the inherent ability to cross the brain endothelial layer, yet at a lower transcytosis rate. Our findings of the interactions driven by HA and Tf on sEVs transport across BECs in vitro and in vivo may serve to design more effective drug delivery strategies for a variety of neurological disorders, including neurodegenerative, inflammatory and infectious diseases.

## ACKNOWLEDGMENT

The authors would like to acknowledge the funding from the European Commission through the Marie Skłodowska-Curie Innovative Training Network “NANOSTEM” (ref. 764958) and EC pathfinder open REGENERAR (Ref: 101129812); the Portuguese funding research institution (FCT) for the funding through the projects EXPL/BTM-ORG/1348/2021, 2022.07615.PTDC and 2022.02803.PTDC; and the PRR project HfPT- Health from Portugal (ref. 02/C05-i01.01/2022.PC644937233-00000047). IA would like also to acknowledge the FCT fellowship 2021.07471.BD. ML acknowledge to FCT for the funding of CEEC project 2021.03756.CEECIND/CP1656/CT0016 (DOI: 10.54499/2021.03756.CEECIND/CP1656/CT0016). The authors would like to thank the Advanced BioImaging and BioOptics Experimental Platform (Champalimaud Foundation, Lisbon) for the assistance in image acquisition and processing in the Zeiss Observer 7, the Bioimaging Unit (IMM, Lisbon) for the assistance with super-resolution imaging in the Zeiss LSM980 Airyscan 2, the Advanced Light Microscopy scientific platform (i3S, Porto) for assistance with STED microscopy, and the Biochemical and Biophysical Technologies facility (i3S, Porto) for the assistance in the SPR experiments and data analysis.

## Author Contributions

IA designed and performed the experiments, analyzed and interpreted the data, and wrote the manuscript. EA and AT co-designed and assisted with SPR experiments. PS assisted with STED microscopy experiments. ML and LF designed and supervised the research. All authors have given approval to the final version of the manuscript.

## Supporting Information

Characterization of native and engineered EVs, Synthesis of acrylated hyaluronic acid, Characterization of the hCMEC/D3 cell line, Representative SPR sensorgrams, Cell viability assayed by hoescht/PI staining, Transferrin and HSA co-localization with the recycling endosome marker Rab11, EVs co-localization with the recycling endosome marker Rab11, Size distribution profile of Cy7-labelled EVs and EVs-HA-Tf, EVs biodistribution ex vivo, Transferrin receptor (TfR) expression in the brain.

## ABBREVIATIONS

BBB: Blood-brain barrier
BEC: Brain endothelial cell
BSA: Bovine serum albumin
CNS: Central nervous system
CSF: Cerebrospinal fluid
HA: Hyaluronic acid
HAS: Human serum albumin
JAM: Junction adhesion molecule
LDLR: Low-density lipoprotein receptor
MISEV: Minimal Information for Studies of Extracellular Vesicles
NTA: Nanoparticle tracking analysis
RMT: Receptor-mediated transcytosis
SEM: Standard error of the mean
sEV: Small extracellular vesicle
SNARE: Soluble NSF attachment protein receptor
SPR: Surface-plasmon resonance
STED: Stimulated emission depletion
TEM: Transmission electron microscopy
Tf: Transferrin
TfR: Transferrin receptor
ZO: Zonula occludens.

