## Supplementary material for "Dual Ligand Cooperation at the Plasma Membrane Drives Transport of Engineered Small Extracellular Vesicles Across Brain Endothelial Cells"

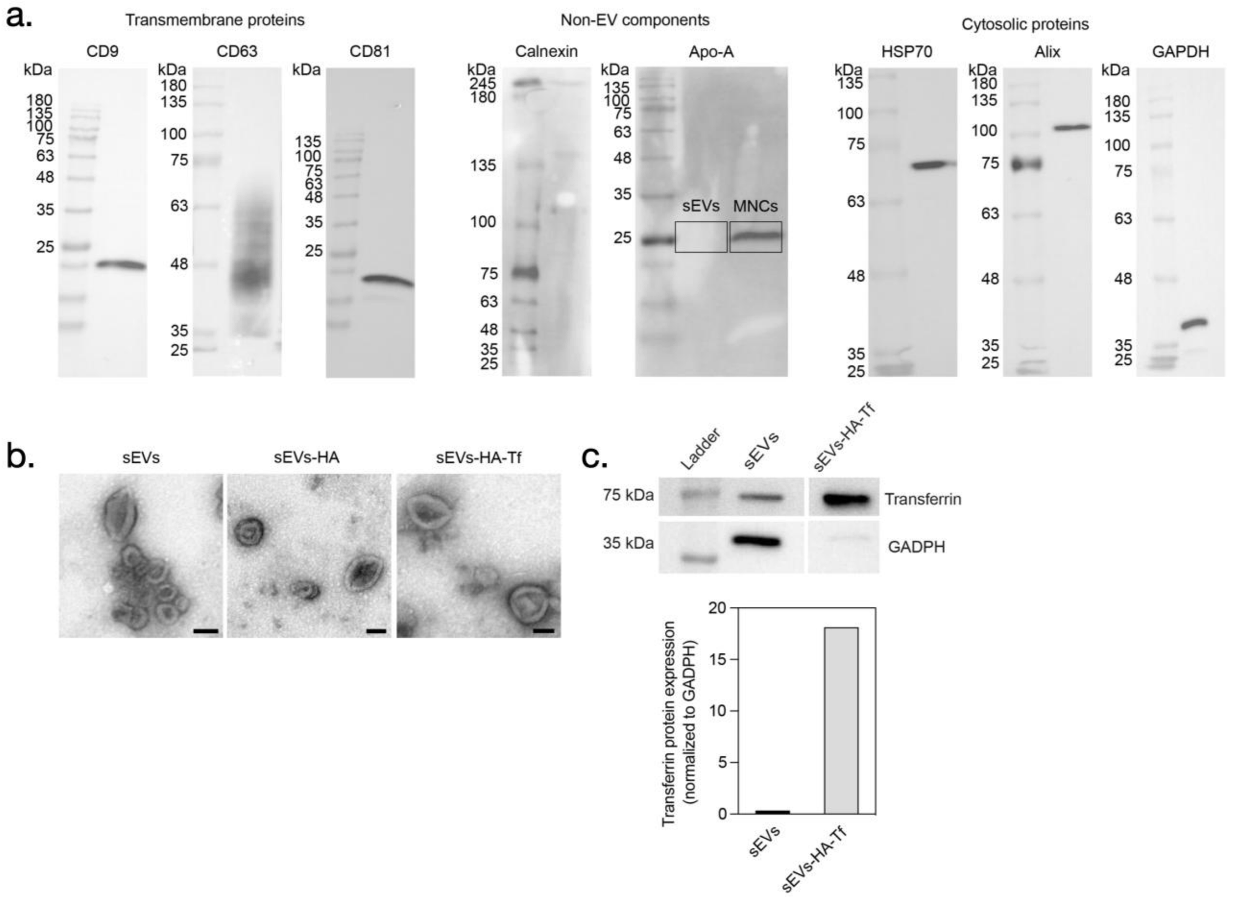


**Supplementary Figure 1.** Characterization of native and engineered sEVs (sEVs-HA and sEVs-HA-Tf). a. Protein-content characterization of human umbilical cord-blood plasma-derived extracellular vesicles by Western Blot. b. Electron-microscopic observation of whole-mounted sEVs. Scale bar: 100 nm. c. Western blot analysis of Transferrin expression in native and engineered sEVs, normalized to GAPDH. A representative Western blot is given plus quantitative analysis of Transferrin band density.

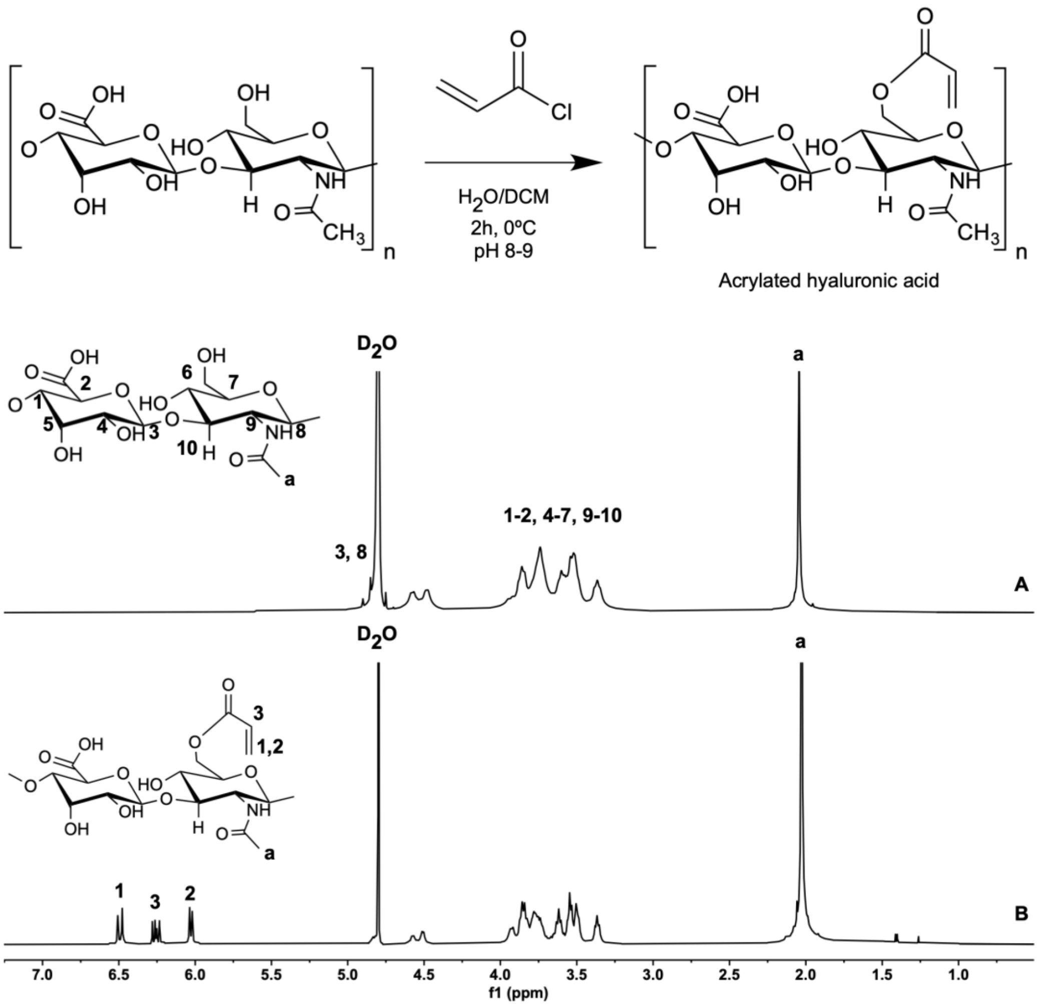


**Supplementary Figure 2.** Synthesis of acrylated hyaluronic acid. Scheme for reaction of HA and acryloyl chloride (AC) and 1H-NMR spectra of (A) bare HA and (B) HA-A (DS= 20%), in D2O, obtained by a 400 MHz spectrometer.

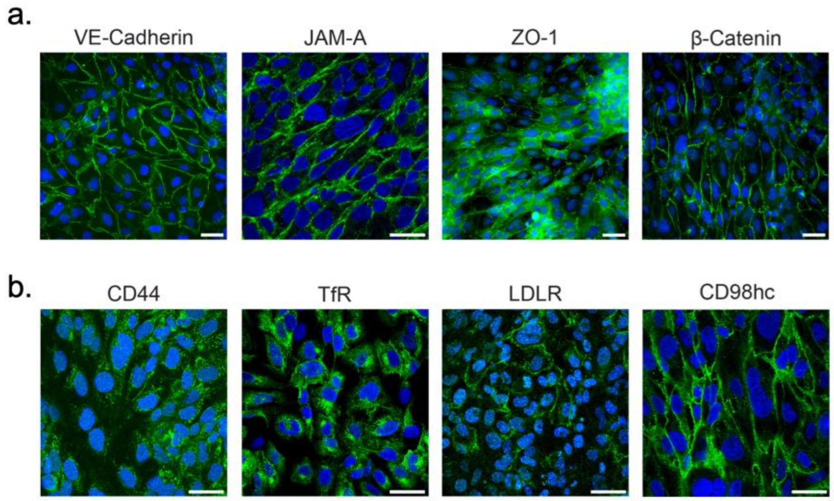


**Supplementary Figure 3.** Characterization of the hCMEC/D3 cell line. a. Confluent monolayers of hCMEC/D3 cells were stained for the endothelial junctional marker VE-Cadherin and for the junction-associated proteins JAM-A, ZO-1 and β-Catenin. b. Expression of the target protein markers, CD44, TfR, LDLR and CD98hc. Scale bar: 40 µm.

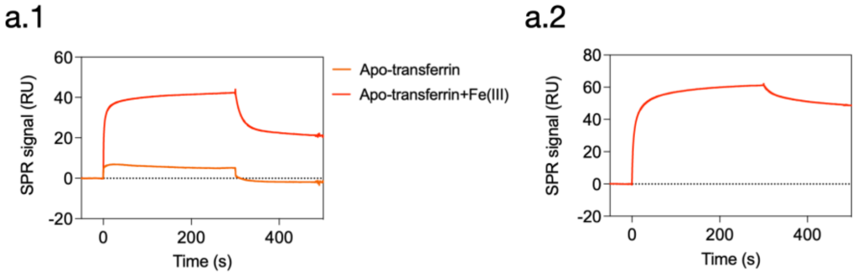


**Supplementary Figure 4.** Representative SPR sensorgrams profiles for a.1. apo-Transferrin, apo-Transferrin pre-loaded with Fe (III) and a.2. holo-Transferrin binding to the immobilized human Transferrin receptor, injected over the surface at the concentration of 400 nM.

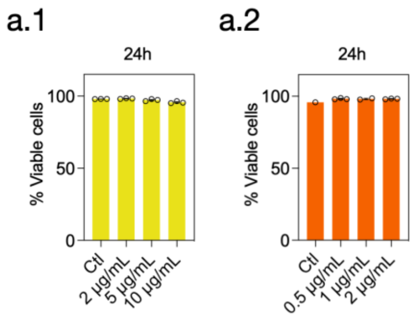


**Supplementary Figure 5.** a.1. Cell viability assayed by hoescht/PI staining for different concentrations of anti-CD44 and a.2. anti-TfR antibodies, over 24 h exposure with the antibodies.

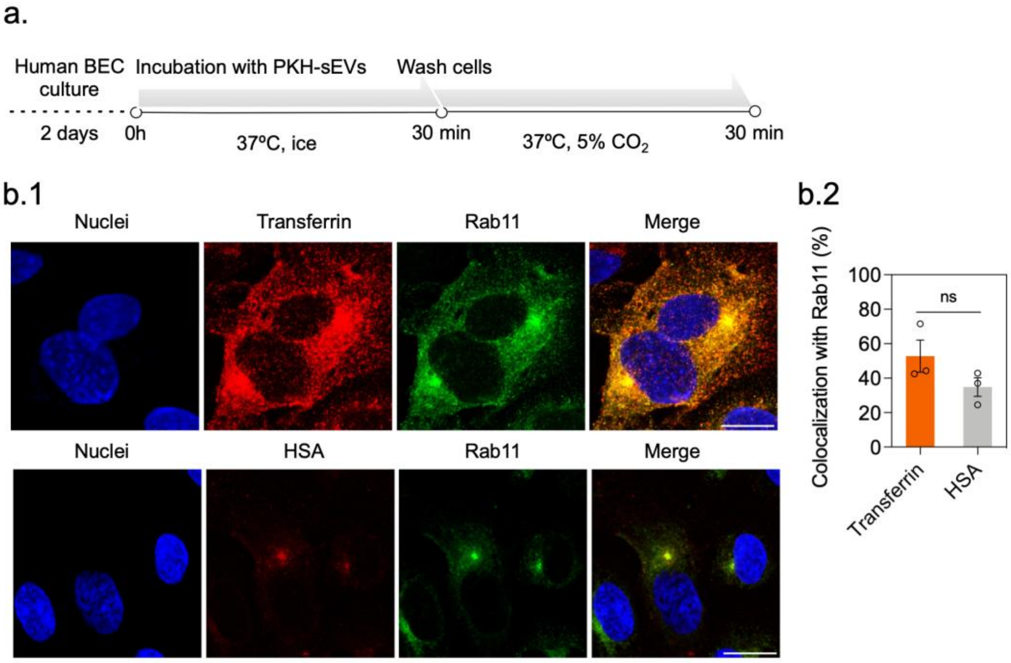


**Supplementary Figure 6.** Transferrin and HSA co-localization with the recycling endosome marker Rab11. a. Scheme of the Transferrin trafficking assay. b.1. Co-localization of the control proteins, Transferrin and HSA with Rab11. Images were acquired by fluorescence confocal microscopy (LSM 710, Zeiss) with a 100x objective. Scale bar: 40 µm. b.2. Co-localization with Rab11 given by Manders' coefficient M1 represent the overlap between Transferrin and HSA with Rab11 positive compartments. Results are expressed as mean±SEM (n=3 independent experiments). Statistical analysis was performed by an unpaired t-test ($p<0.05$).

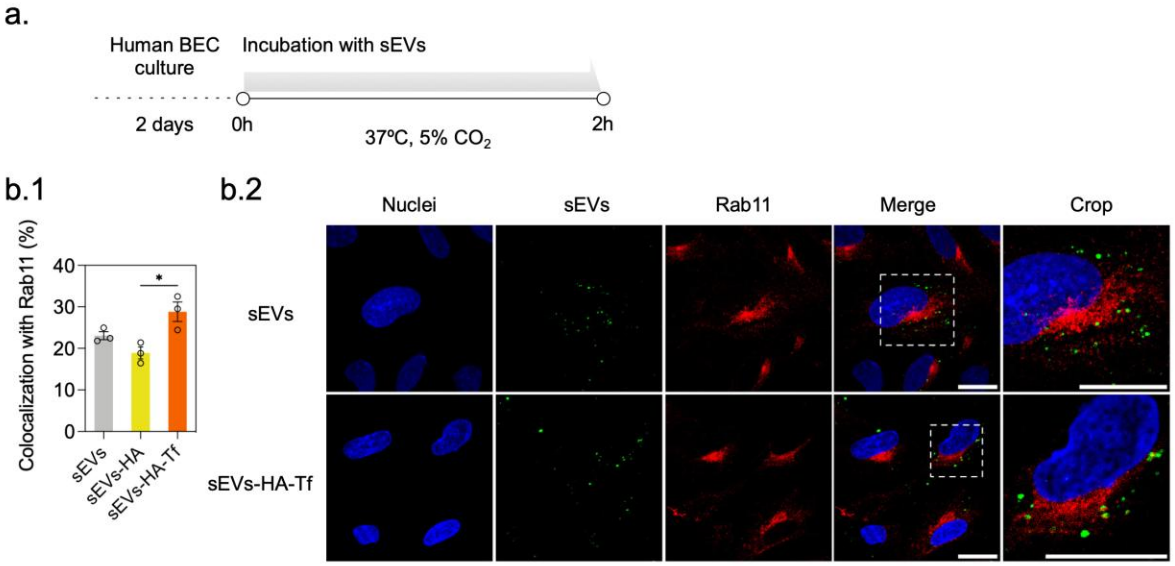


**Supplementary Figure 7.** sEVs co-localization with the recycling endosome marker Rab11. a. Schematic drawing of the sEVs trafficking assay. b.1. Co-localization with Rab11 given by Manders' coefficient M1 representing the overlap between sEVs and Rab11 positive compartments. Results are expressed as mean±SEM (n=3 independent experiments). Statistical analysis was performed by an unpaired t-test ($p<0.05$). b.2. Overview of sEVs and sEVs-HA-Tf co-localization with Rab11. Images were acquired by fluorescence confocal microscopy (LSM710, Zeiss) with a 100x objective. Scale bar: 40 µm.

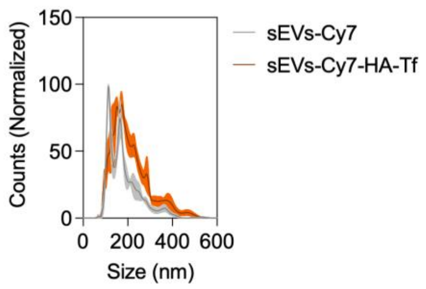


**Supplementary Figure 8.** Size distribution profile of Cy7-labelled sEVs and sEVs-HA-Tf evaluated by NTA, expressed in particle counts (percentage) as a function of particle size diameter (nm). Results are mean ± SEM (3 independent experiments). The standard error of the mean is represented in the color filling.

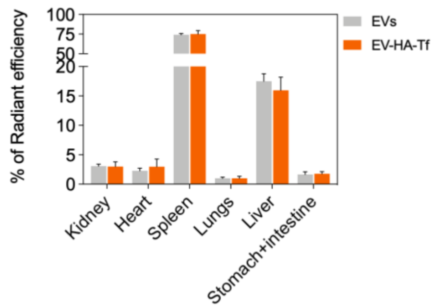


**Supplementary Figure 9.** sEVs biodistribution ex vivo, in the different peripheral organs 1 h after systemic administration. Results are expressed as mean ± SEM (n=5 animals per experimental condition).

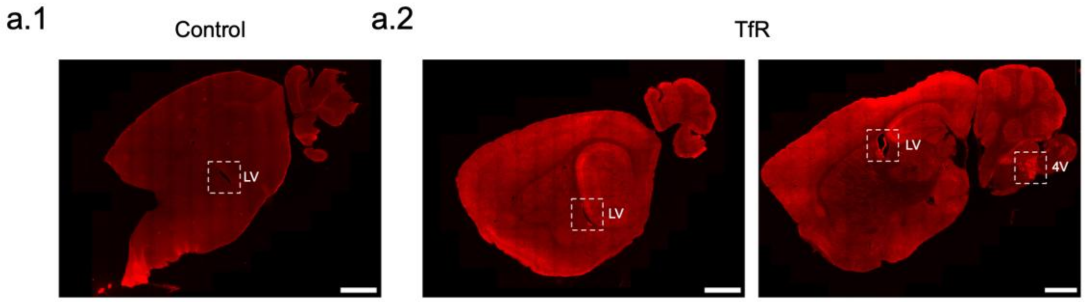


**Supplementary Figure 10.** Transferrin receptor (TfR) expression in the brain. a.1. Sagittal section showing the control, in the absence of primary antibody, and a.2. TfR differential expression in the lateral (LV) and fourth ventricles (4V). Images were acquired in a slide scanner (AxioScan7, Zeiss) with a 20x objective. Scale bar: 1 mm.
